# Seqpac: A New Framework for small RNA analysis in R using Sequence-Based Counts

**DOI:** 10.1101/2021.03.19.436151

**Authors:** Signe Skog, Lovisa Örkenby, Unn Kugelberg, Kanwal Tariq, Ann-Kristin Östlund Farrants, Anita Öst, Daniel Nätt

## Abstract

Small RNA sequencing (sRNA-seq) has become important for studying regulatory mechanisms in many cellular processes. Data analysis remains challenging, mainly because each class of sRNA—such as miRNA, piRNA, tRNA- and rRNA-derived fragments (tRFs/rRFs)—needs special considerations. Analysis therefore involves complex workflows across multiple programming languages, which can produce research bottlenecks and transparency issues. To make analysis of sRNA more accessible and transparent we present seqpac: a tool for advanced group-based analysis of sRNA completely integrated in R. This opens advanced sRNA analysis for Windows users—from adaptor trimming to visualization. Seqpac provides a framework of functions for analyzing a PAC object, which contains 3 standardized tables: sample phenotypic information (P), sequence annotations (A), and a counts table with unique sequences across the experiment (C). By applying a sequence-based counting strategy that maintains the integrity of the fastq sequence, seqpac increases flexibility and transparency compared to other workflows. It also contains an innovative targeting system allowing sequence counts to be summarized and visualized across sample groups and sequence classifications. Reanalyzing published data, we show that seqpac’s fastq trimming performs equal to standard software outside R and demonstrate how sequence-based counting detects previously unreported bias. Applying seqpac to new experimental data, we discovered a novel rRF that was down-regulated by RNA pol I inhibition (anticancer treatment), and up-regulated in previously published data from tumor positive patients. Seqpac is available on github (https://github.com/Danis102/seqpac), runs on multiple platforms (Windows/Linux/Mac), and is provided with a step-by-step vignette on how to analyze sRNA-seq data.

## BACKGROUND

The past decades have uncovered a diversity of small RNA (sRNA), which differs greatly in their biogenesis and biological roles. This involves miRNA that is generated from transcribed precursors and recruited by Argonaute proteins for post- and pre-transcriptional gene silencing (1–5). Having a similar mechanism, piRNA primarily silence repetitive transposable elements in the germline, and can be amplified by means of the so-called ping-pong cycle (6). Other classes involves rRNA and tRNA derived fragments (rRF/tRFs) that may interact with Argonaute proteins in a piRNA/miRNA-like fashion, but may also directly interfere with translational processes in the ribosome (7–10). Some tRFs may not even align to their genome of origin, since their parental tRNA matures post-transcriptionally by receiving additional nucleotides (11). While many sRNA classes exerts their function in the cytoplasm, some intermediately sized none-coding RNA—like the snoRNA, scaRNA and snRNA—are associated with specific organelles inside the nucleus where they play important roles in the post-transcriptional shaping (splice, fold, and modify) of other RNA molecules (12, 13).

This complexity, where some sRNA may target single gene products while others target highly repetitive regions, where some are biologically active after transcription while others are post-transcriptionally modified prior to activation, where some align to the genome that they originated from while others do not, makes the analysis sRNA challenging. Today, it is also becoming increasingly popular to apply high-throughput sequencing in sRNA experiments, which makes the analysis even more complicated. Combining massively parallel sequencing with specialized library preparation protocols that select for short RNA species generate data often containing millions of unique short RNA sequences across tens-to-hundreds of samples.

Several tools and pipelines, such as Sports (14), MintMap (11), sRNAtoolbox (15), sRNAnalyzer(16), COMPSRA (17), and iSmaRT (18) have been developed to overcome some of the analytical thresholds in sRNA analysis. As a rule, these tools wrap around multiple programs written in multiple programming languages, such as cutadapt (19) for adapter trimming, bowtie (20) for genome mapping, and subread:featureCounts (21) for counting sequences across sRNA subspecies. Thus, in labs that lack strong programming skills and previous experience of sRNA analysis, troubleshooting and advanced analysis often become bottlenecks. This may result in the exclusion of ‘difficult-to-analyze’ sRNA in favor of more straight-forward sub-species, such as miRNA. Unless better, more coherent, and user-friendly tools are developed, such discrimination will result in severe literature biases.

Workflows for sRNA-seq analysis regularly build on methods from gene-centric DNA/RNA-seq approaches, such as regular mRNA-seq. This usually involves mapping individual samples against a reference genome followed by counting overlaps of genomic coordinates between sample reads and known genomic features, such as gene exons or miRNAs. Such feature-based counting (Figure 1A) is often done one read and one sample at the time. Most sRNA experiments, however, do not contain a single sample. Instead, they contain multi-sample groups. Therefore, as an alternative, read sequences across the whole experiment can be counted prior to aligning the read to a reference genome. Such sequence-based counting (Figure 1A) would prevent annotating the same sequence multiple times both within and across samples. More importantly, this strategy would maintain sequence integrity. Thus, further annotation of the counted sequences would be possible at any time during the analysis. In addition, with sequence-based counting users may choose to remove sequences with low evidence, which fails to replicate across their experiment. Hypothetically, these advantages with sequence-based counting may not only have dramatic effects on computational performance. It may also increase the transparency and flexibility of the whole analysis.

**Figure 1.**
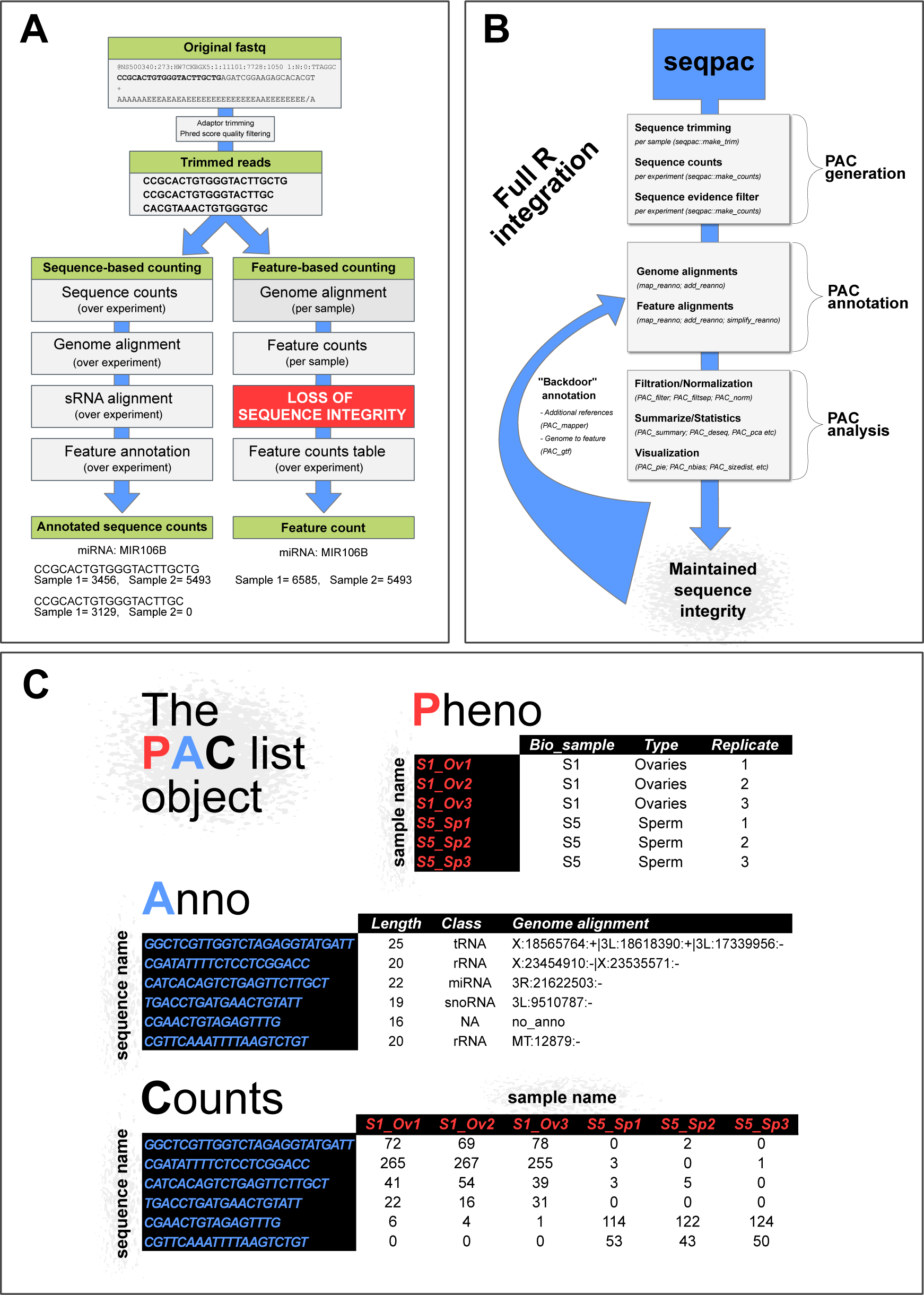
Small RNA sequence analysis using seqpac. (A) The difference between sequence-based counting and feature-based counting in sRNA analysis. With sequence-based counting an experiment-wide count table can be created before genome alignment. This allows for efficient mapping and for sequence integrity to be maintained through the analysis. In contrast, feature-based counting strategies counts overlaps between reads’ genome alignment and coordinates of genomic features. This is less efficient and disrupts sequence integrity. (B) Sequence-based counting is central in the seqpac workflow, which is completely integrated in R, from fastq adaptor trimming and preprocessing to group-based visualization and statistical analysis. (C) Secpac builds a framework of functions that processes and analyzes a standardized list: the PAC object. In its simplest form PAC contains three tables: the Pheno table with sample information; the Anno table with sequence information, and the Counts table containing the counts of sequences across samples.

Here, we present—seqpac—a novel framework for sequence-based multi-sample sRNA analysis. From adapter trimming to the visualization of group-differences, seqpac is completely integrated as an open-source package in R. This makes it accessible from multiple platforms, including Windows, Mac and Linux. Using both published and novel data, we show that sequence-based counting combined with a multi-sample approach, not only positively affects computational performance, making sRNA-seq analysis accessible on a standard computer. It also increases the flexibility and transparency throughout the analysis. We illustrate this by detecting severe contamination in published data that was previously analyzed using a feature-based counting strategy. Finally, we use the strengths of seqpac to discover and confirm a novel rRF implicated as a diagnostic/prognostic marker in cancer.

## MATERIALS AND METHODS

### 1. Package development

Seqpac is available for download at github (https://github.com/Danis102/seqpac). Procedures on how to install seqpac are explained in the vignette (https://github.com/Danis102/seqpac/tree/master/vignettes). Dependencies for the main seqpac functions are listed in Table 1. Seqpac was developed and tested on a Linux Mint v.19.1 computer using R 3.4.4 in RStudio 1.2.1335 and devtools 2.3.2. The computer had an Intel Core i7-9800X CPU at 3.8 GHz (8 cores with in total 16 threads) and contained 94 Gb of ram memory. All R internal functions (e.g. make_cutadapt excluded) were subsequently tested on multiple Windows 10 computers using R 3.6.3 and 4.0.1.

**Table 1.** Quick reference for seqpac

| Family | Description | Function name | Dependence | Function description |
| --- | --- | --- | --- | --- |
| **PAC generation** | Prepares and builds a PAC object from fastq files. | *make_counts* | *foreach, ShortReads* | Reads fastq, calls *make_trim* or *make_cutadapt*, performs low-level filter, and counts sequences. |
|  |  | *make_trim* | *foreach, Biostrings* | Adaptor trimming and fastq quality filter using R internal packages |
|  |  | *make_cutadapt* | *foreach, cutadapt, fastq_quality_filter* | Adaptor trimming and fastq quality filter using external installed software |
|  |  | *make_anno* | - | Generates a simple annotation table from a count table. |
|  |  | *make_pheno* | - | Prepares user provided phenotype table containing sample information. |
|  |  | *make_PAC* | - | Builds PAC object with Pheno, Anno and Counts tables. |
|  |  | *PAC_check* | - | Checks if a PAC object is compatible with seqpac functions |
| **PAC annotation** | Sequence annotations by aligning against fasta references or overlap with genomic coordinates. | *map_reanno* | *Rbowtie/bowtie* | Reannotates PAC sequences through progressive mismatch cycles. |
|  |  | *import_reanno* | *foreach, tibble* | Called by *map_reanno* to import bowtie output into R. |
|  |  | *add_reanno* | *foreach, tibble* | Builds a reannotation object with genome coordinates and/or classified fasta sequence names. |
|  |  | *simplify_reanno* | - | Makes hierarchical classifications from classified fasta sequence names. |
|  |  | *PAC_gtf* | *tibble, rtracklayer, GenomicRanges* | Overlaps genomic coordinates of PAC sequences with features of a gtf file. |
|  |  | *PAC_mapper* |  | Backdoor to the reanno workflow for fast mapping of PAC resulting in a map object. |
|  |  | *map_rangetype* |  | Classifies sequences in map objects according to ranges or secondary structures (e.g. 5', i', 3') |
| **PAC analysis (preprocessing)** | Filtering and normalization. | *PAC_filter* | - | Subsets data by target objects or coverage thresholds. |
|  |  | *PAC_filtersep* | - | Extracts sequences reaching a threshold within groups of a pheno_target object. |
| | | *PAC_norm* | *DESeq2* | Normalize a raw counts table and saves it in the PAC\$norm 'folder'. |
| **PAC analysis (statistics)** | Performs statistical analyses and visualizations. | *PAC_summary* | - | Simple summaries using pheno_targets (means, sd, se, %diff, log2fc) saved in PAC\$summary. |
|  |  | *PAC_deseq* | *foreach, DESeq2* | Prepares, performs and plots DESeq2 analysis from PAC object. |
|  |  | *PAC_pca* | *FactoMineR, factoextra* | Performs principal component analysis and plots the results. |
|  |  | *PAC_saturation* | *foreach, ggplot2* | Performs a sequence saturation analysis and plots the results. |
| **PAC analysis (visualization)** | Generates graphs and saves processed data summarized over both phenotype and annotations. | *PAC_pie* | *ggplot2, cowplot, grDevices* | Pie-plots using pheno_target and anno_target. |
|  |  | *PAC_stackbar* | *ggplot2, ggthemes, reshape2, grDevices* | Stacked bar diagrams using pheno_target and anno_target. |
|  |  | *PAC_jitter* | *ggplot2* | Jitter plots using pheno_target and anno_target. |
|  |  | *PAC_nbias* | *ggplot2, ggthemes, grDevices* | Size distributed histogram stacked by nucleotide at a defined position (e.g. 1st nucleotide bias). |
|  |  | *PAC_sizedist* | *ggplot2, ggthemes, grDevices* | Size distributed bars stacked by an anno_target column (e.g. miRNA/piRNA size distributions). |
|  |  | *PAC_covplot* | *ggplot2, reshape2, grDevices, GenomicRanges* | Plots PAC sequence coverage over a reference sequence such as a tRNA or rRNA. |
|  |  | *PAC_trna* | *ggplot2, reshape2, grDevices* | tRNA fragment analysis using information from range-classified map object. |

All functions are described in detail in the manual for each function (e.g. '?make\_counts' in the R terminal) and are exemplified in the seqpac vignette; 'vignette("seqpac")' in R terminal).

### 2. Testing seqpac using published datasets

Fastq files for 4 datasets were accessed through Sequence Reads Archive (SRA) and European Reads Archive (ENA). We prefer downloading these files and their metadata through ENA (https://www.ebi.ac.uk/ena). All code for processing and generating the results presented in Figure 2, 4, 7 and 8 are available in Supplementary text S1. A brief explanation is provided below.

**Figure 2.**
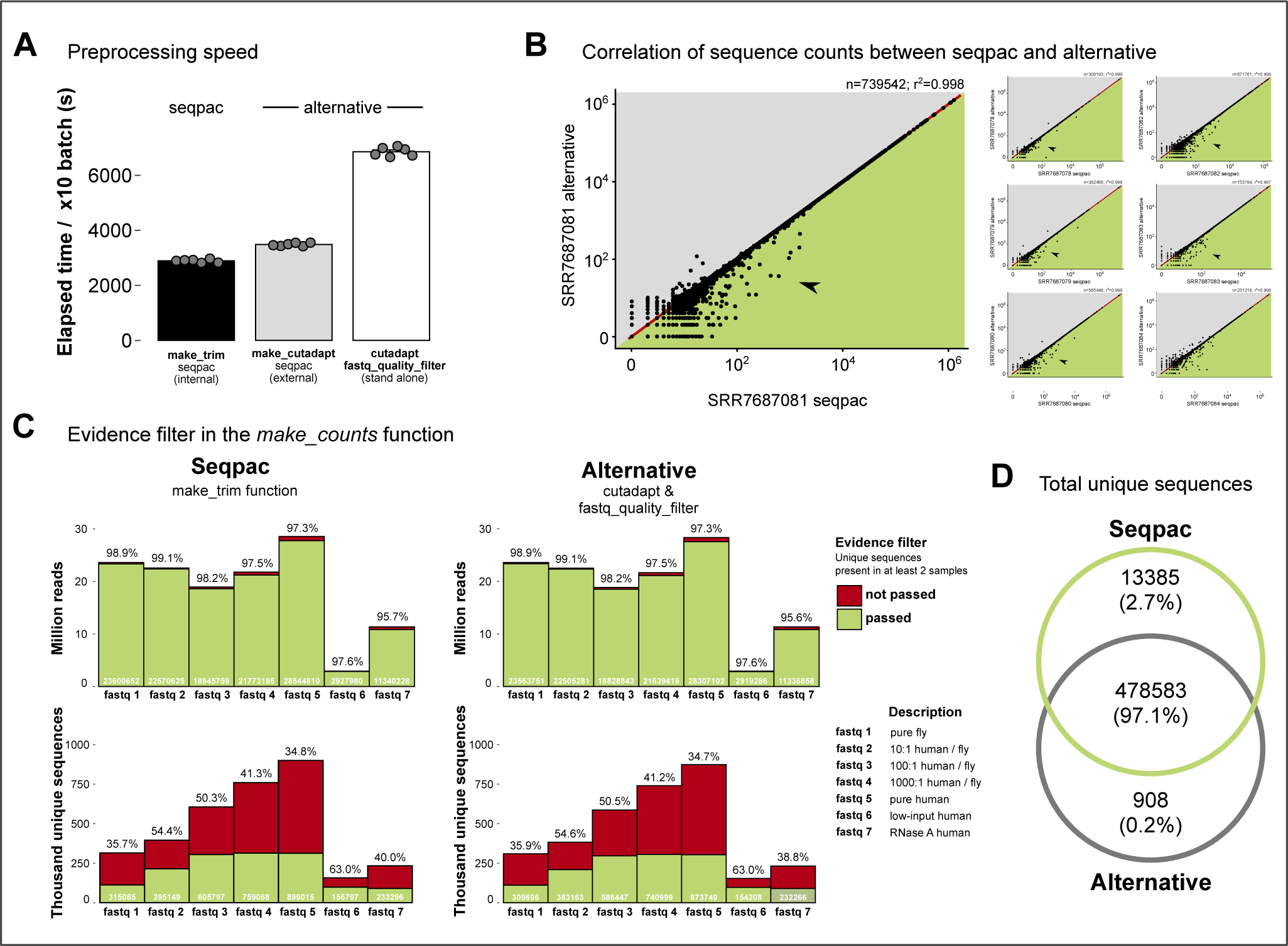
Performance of sepac’s internal trimming and counting functions. Seqpac contains multiple options for trimming and filtering adaptor sequences prior to generating a count table. The make_counts function counts sequences of already trimmed fastq files or calls make_trim (R internal) or make_cutadapt (R external) functions prior to counting. Using the Kang et al. 2018 dataset (SRA access: PRJNA485638), (A) shows side-by-side the preprocessing time for seqpac’s trimming functions and a popular alternative workflow based on the cutadapt and fastq_quality_filter functions. The test involved 7 fastq files iterated 10 times over 6 batches per function using 7 parallel processes. (B-D) Further evaluated performance of make_trim in terms of the output dataset. (B) While sequence counts strongly correlated with the alternative workflow, make_trim more often generated higher counts (arrows). This was primarily a result of concatemer trimming in make_trim (see main text). (C) The make_counts function also contains an evidence filter, which in default mode discard sequences that fails to replicate in at least two independent fastq files. Normally, this low-level filter maintains most reads (top bars), while limiting the sequence diversity (bottom bars). While make_trim and the alternative generated very similar datasets after evidence filters, make_trim generated slightly more unique sequences, which was confirmed by Venn-diagram (D) showing higher ratio of sequences unique to the make_trim workflow.

**Figure 3.**
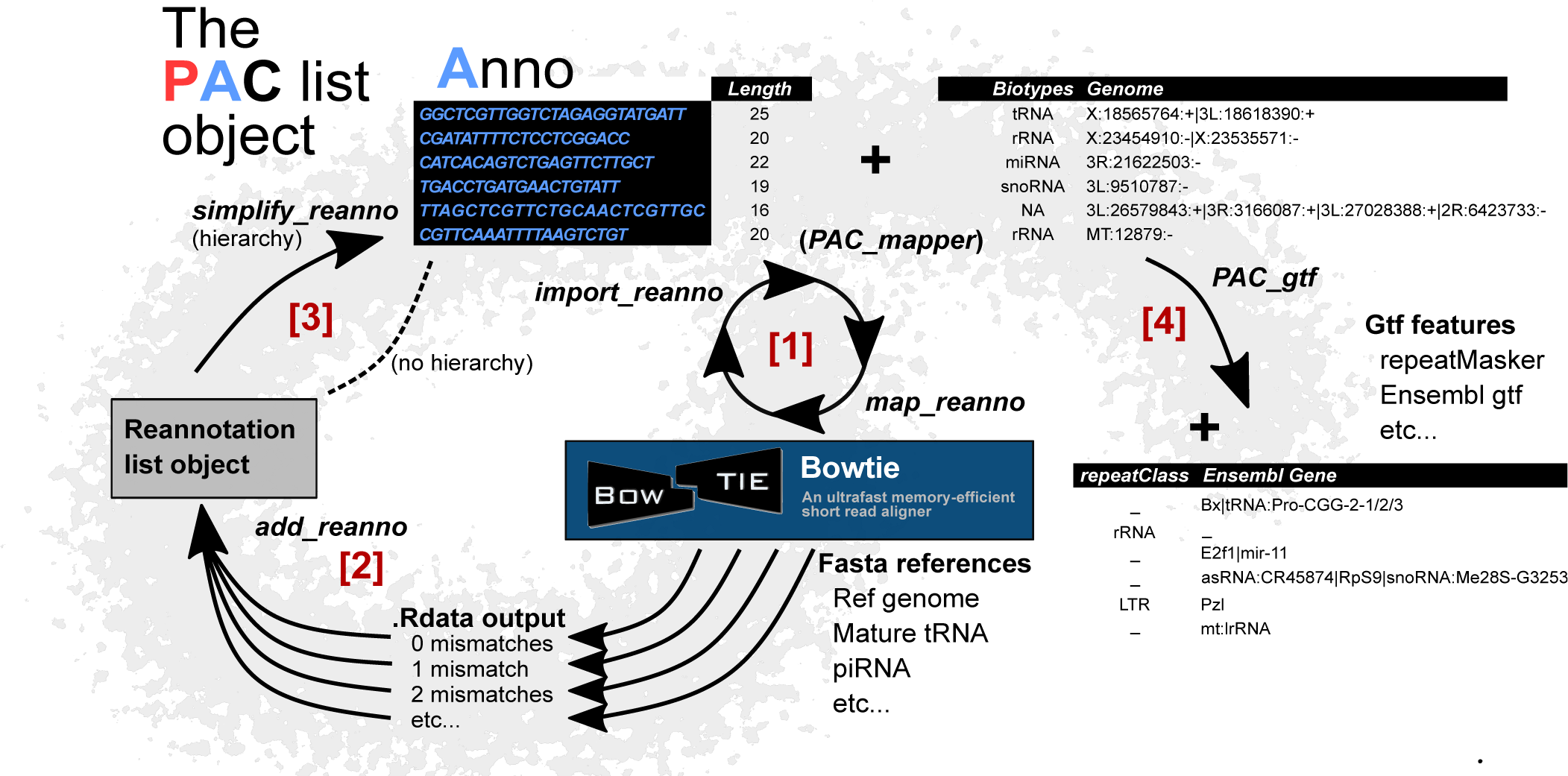
Annotating read sequences in PAC objects. For annotation of read sequences, seqpac mainly relies on the re-annotation workflow [1-3]. The *map_reanno* and *import_reanno* functions use Bowtie to align PAC sequences against references sequences, e.g. species genome or sRNA database [1]. This is done over cycles where each cycle introduces 1 additional mismatch in the mapping, and where only read sequences with no alignment proceed to the next cycle. After the mismatch cycles, *add_reanno* reads the resulting .Rdata files and organize the output into a reannotation list object [2]. Tables in this list can either directly be merged with a PAC annotation table or can be simplified hierarchically using the *simplify_reanno* function [3]. The *PAC_mapper* function is a convenient wrapper for smaller reference sequences (e.g. tRNAs or rRNAs) that will automatically generate Bowtie indexes. [4] After a PAC object has been aligned to a genome, the *PAC_gtf* can be used to overlap genomic coordinates of PAC sequences with known coordinates for genomic features, e.g. repeats and protein coding exons.

#### 2.1 Kang et al. 2018 – Benchmarking and reannotation using human and fruit fly multi-genome samples (Figure 2, 4)

Kang et al. 2018 (22) (SRA accession: PRJNA485638; ENA download: https://www.ebi.ac.uk/ena/browser/view/PRJNA485638) were used for benchmarking seqpac’s make_trim function against two similar workflows. In both alternative workflows, system calls to cutadapt (19) and fastq_quality_filter (in FASTX-Toolkit; http://hannonlab.cshl.edu/fastx_toolkit/) were made from within R. The first used the make_cutadapt function to replicate the parallelization for make_trim using the foreach package (23), while the second used the internal parallelization option in cutadapt. System time was monitored over 10 iterations replicated 6 times using the rbenchmark package (24).

PAC objects with counts from trimming/filtering using the *make_trim* function and cutadapt/fasq_quality_filter alternative, were generated using *make_counts* function either with *trimming=”seqpac” or trimming=”cutadapt”*. Bar graphs from the low-level evidence filtering were saved. To assure that only sRNA were include, since this dataset was generated from a 75 cycle flow-cell, we removed reads that failed to contain adaptor sequence and only kept reads <=45 nt. The counts lists with progress reports were then applied to the standard PAC generation workflow (*make_counts > make_anno > make_pheno > make_PAC*). As phenotypic input file for *make_pheno* function we used metadata downloaded from SRA/ENA.

After benchmarking, only the internal (*make_trim*) PAC object was applied to the reannotation workflow. Reannotation against either the human and fly reference genomes or sRNA class references were applied, using either the *map_reanno* import=”genome” or import=”biotype” options, respectively. For genome alignments we downloaded Homo sapiens GRCh38.101 (hg38) and Drosophila melanogaster BDGP6.28 (dm6) in fasta references at Ensembl ftp (http://www.ensembl.org/info/data/ftp/). For the sRNA class alignment we downloaded fasta references for miRNA (mirBase v.21), ncRNA (Ensembl.ncrna), tRNA (GtRNAdb) and piRNA (pirBase) for the human and fruit fly genomes, respectively.

After generating reanno objects in R using the *make_reanno* function, we added and simplified the annotations using the *add_reanno* and *simplify_reanno* functions. The sRNA class hierarchy in *simplify_reanno* was set to rRNA > tRNA > miRNA > snoRNA > snRNA > lnc/lincRNA > piRNA. Plots in Figure 4 were generated using the *PAC_pie, PAC_sizedist* and *PAC_nbias* functions.

**Figure 4.**
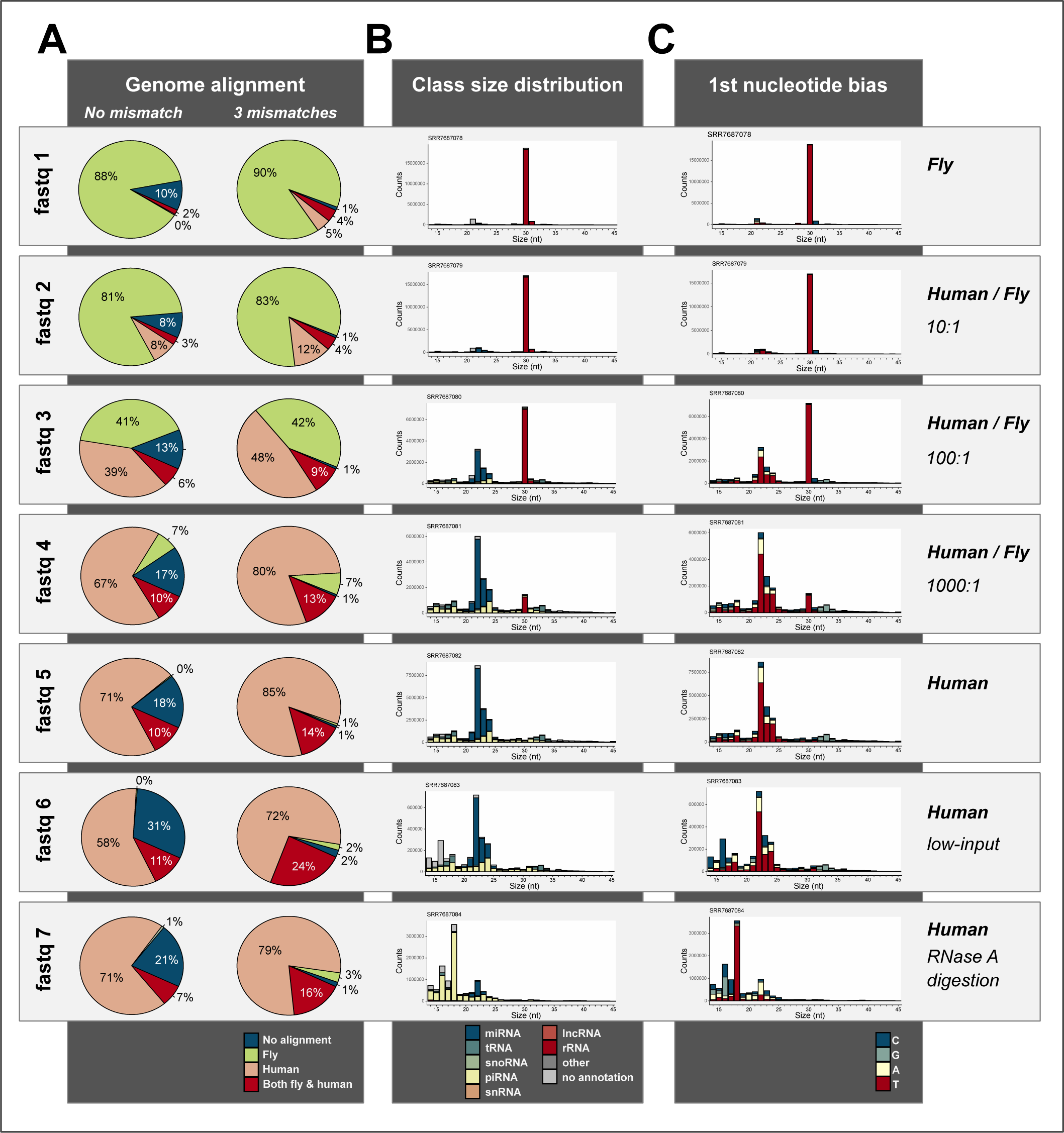
Multi-genome sRNA analysis demonstrating the strength in seqpac reannotation workflow. Graphs were plotted using a PAC object generated from the Kang et al. 2018 dataset (SRA access: PRJNA485638) primarily developed for studying interspecies contaminated samples where RNA from fly (S2) and human (HEK-293T) cells was mixed in different ratios. Seqpac functions used for generating the graphs were: (A) *PAC_pie* for genome proportion pie charts, (B) *PAC_sizedist* for size distribution histograms with sRNA class annotation, and (C) *PAC_nbias* for the frequency of the first nucleotide stratified over the sequence size distribution.

#### 2.2 Tong et al. 2020 – Detecting contamination in cancer cell lines (Figure 7, 8)

Tong et al. from 2020 (25) (SRA accession: PRJNA666144; ENA download: https://www.ebi.ac.uk/ena/browser/view/PRJNA666144) were used for exemplifying the strengths of sequence-based counting in detecting severe bias in cancer cell line experiements. PAC generation and reannotation was performed similarly to the Kang et al. dataset (MATERIALS AND METHODS 2.1) with a few exceptions. Since the Tong et al. was generated from a 50 cycle flow-cell we did not remove reads that failed to contain adaptor sequence and did not filter by max read length. Analysis and graphs were generated using the *PAC_pca, PAC_sizedist* and *PAC_stackbar* functions. We also verified the 5’ ETS rRF of the 45S pre-rRNA (NR_146144.1) with the *PAC_mapper* and *PAC_covplot* functions using the same fasta reference as described in Methods 2.3. When reannotating the PAC object after the initial analysis, we used the Mycoplasma hyorhinis ATCC (ASM38351v1) genome in parallel with Homo sapiens (hg38).

#### 2.3 Skog et al. 2021 – HeLa anti-cancer treatment dataset (current study; Figure 8)

The anti-cancer treatment dataset was generated in the current study (see Methods 3) and is available at SRA (accession: PRJNA708219). Since this dataset was generated from a 75 cycle flow-cell, an annotated PAC object was created as for the Kang et al. dataset (see Methods 2.1) removing reads that failed to contain adaptor sequence. To better compare with the Tong et al. dataset we set a max read length of 65 nt. The *PAC_deseq* function was used to initially identify BMH21 sensitive fragments comparing cells exposed to BMH21 for 60 min to those exposed to DMSO for 60 min (control). Mapping against pre-rRNA was done using the *PAC_mapper* function with a custom fasta reference (Supplementary file S2). This reference first contained the GenBank sequence NR_146144.1. After identifying 4 peaks using the *PAC_covplot* function, we added the zoomed in regions of chr 21 aligning with NR_146144.1 and containing each of the four rRF peaks (Peak 1 = chr21:8206319-8206669, Peak 2 = chr21:8212475-8212825, Peak 3 = chr21:8213765-8214115, Peak 4 = chr21:8218787-8219137). These regions were downloaded from the UCSC genome browser. Finally, we added the 47S GenBank entry U13369.1 to the fasta.

#### 2.4 Xu et al. 2020 – Validation in cervical cancer patients (Figure 8)

The Xu et al. 2020 (26, 27) (SRA accession PRJNA607023; ENA download: https://www.ebi.ac.uk/ena/browser/view/PRJNA607023) dataset, used for validating the 5’ ETS rRF of the 45S pre-rRNA (NR_146144.1) in clinical samples. We only used the 8 fastq files obtained by sRNA size-fractions. Files were generated using a paired-end 2x150 cycle flow cell kit. Thus, we discarded the paired—second—read and only kept the trimmed sequences of the first read where an adaptor was present (as in see Methods 2.1 and 2.3).

## 3. Generating the HeLa anti-cancer treatment dataset

Adherent HeLa CLL-2 cells were obtained from ATCC and were maintained at 37°C and 5% CO2 in high glucose Dulbecco’s modified Eagle’s medium (DMEM), supplemented with 10% fetal bovine serum and 1% Penicillin/Streptomycin cocktail. Cells were treated with 1uM BMH-21 in antibiotic free media for 60min and 12h. Cells treated with DMSO for 60 min were used as control. The media was removed, cells were washed with PBS, collected with trypsinization, and stored at -70°C until further processing.

Frozen cells were homogenized in prechilled Qiazol (Qiagen, Hilden, Germany) using a Tissue Lyser LT (Qiagen) set to 2 min at 30 oscillations/second with 5 mm Stainless Steel Beads (Qiagen). RNA was then extracted using miRNeasy Micro kit (Qiagen), and the integrity of purified RNA was confirmed on a Bioanalyzer (Agilent Technologies, Santa Clara, USA), where sample RIN values ranged between 9.3-10. Library preparation was done with NEBNext Small RNA Library Prep Set for Illumina (New England Biolabs, Ipswich, USA) with 100 ng of input total RNA according to manufacturer instructions, except for the following minor customizations: reactions were scaled-down to half the volume, adapters were diluted 1:2, amplification was done for 12 cycles, and libraires were size-selected for 130 to 190 nt fragments on a pre-casted 6% polyacrylamide Novex TBE gel (Invitrogen, Waltham, USA). Gel extraction was done using Gel breaker tubes (IST Engineering, Milpitas, USA) in the buffer provided in the NEBNext kit. After precipitation, the library concentrations were estimated using QuantiFluor ONE ds DNAsystem on a Quantus fluorometer (Promega, Madison, USA). Pooled libraries were sequenced on NextSeq 500 with NextSeq 500/550 High Output Kit version 2.5, 75 cycles (Illumina, San Diego, USA). All pooled libraries passed Illumina’s default quality control.

## RESULTS AND DESCRIPTION

### 1.1 The seqpac workflow

Seqpac comes with a vignette that contains a step-by-step in-depth guide on how to analyze sRNA data from high-throughput sequencing. All functions can be tested using a set of down-sampled fastq files, with sRNA data originating from single fruit fly embryos. A quick reference to the main functions in seqpac is available in Table 1. Scripts for generating many of the analysis presented in the figures are available in Supplementary file S1.

The general seqpac workflow involves three separate steps: constructing, annotating and, analyzing a PAC object (Figure 1B). A PAC object is in its simplest form an R list object, listing a phenotype (Pheno) table with sample information, an annotation (Anno) table with information about unique sequences, and a counts (Counts) table with the counts of sequences across samples (Figure 1C). While this setup reminds of many S4 class objects in packages such as limma (28), DESeq2 (29) and minfi (30) etc., we have deliberately made the PAC list a regular S3 object, holding two classifications ‘PAC’ and ‘list’. One reason is that S4 objects are often a source of confusion for beginners in R. Another is that all basic functions for handling lists are directly applicable on the PAC object, making it easy for more advanced users to customize their workflows.

### 2.1 Constructing the PAC object

Building the PAC object starts by generating a counts table. This is primarily done by the *make_counts* function. It uses fastq formatted sequence files to generate a standardized data frame, where each row represents unique sequences in the experiment, while columns represent samples (Figure 1C). This table maintains the framework for all subsequent analysis. The phenotype and annotation tables contain further information about samples (columns in the counts table) and sequences (rows in the counts table). These tables are produced by the *make_pheno* and *make_anno* functions. The phenotype table is provided by the user and can optionally be merged with a progress report from the adaptor trimming and low-level filtering (see Results 2.2). The *make_anno* function prepares a very primitive annotation table that will expand in the reannotation workflow (see Results 3.1-3.4). Finally, *make_PAC* checks the different components and builds the PAC object.

### 2.2 Trimming fastq of adaptor sequence

The *make_counts* function reads raw sequence files in fastq format using the ShortRead package (31), trims the reads of adaptor sequence and filters low-quality and non-replicable reads, prior to counting each unique read sequence across all samples. For adaptor trimming seqpac has an internal and external alternative.

Internally, *make_counts* calls the stand alone *make_trim* function that primarily uses the Biostrings package (32) to efficiently search and remove any adaptor sequence. In addition, sequences with low quality base scores can be filtered. For the external option, *make_counts* is dependent on system calls to externally installed cutadapt (19) and *fastq_quality_filter* (available in FASTX-Toolkit) (33) software.

To test the performance of seqpac’s *make_trim* function, we downloaded fastq files from the Kang et al. study from 2018 (22) (SRA project: SRP157338). This dataset contains 7 fastq ranging between 52.7-492.8 Mb in compressed size (mean=310.1 Mb) and were generated from either human or fruit fly RNA, where some samples were generated by mixing RNA from these species in different ratios. Using the rbenchmark package (24) we trimmed/filtered these files over 10 iterations replicated 6 times for the *make_trim* and *make_cutadapt* functions, as well as stand-alone *cutadapt/fastq_quality_filter* using near-to-identical settings. Each function was given 7 parallel jobs on a Linux desktop computer (for hardware specifications, see Methods). While make_trim and make_cutadapt uses the foreach package (23) to parallelize jobs across processor cores/threads, the stand-alone *cutadapt/fastq_quality_filter* workflow used cutadapt’s internal parallelization option (- p 7). The *make_trim* function was on average 1.2 times faster than *make_cutadapt*, and 2.4 times faster than *cutadapt/fastq_quality_filter* (Figure 2A). On average, *make_trim finished* trimming/filtering all 7 fastq in 4.8 min, *make_cutadapt* in 5.8 min and the stand-alone alternative in 11.4 min. The slow performance of the stand-alone alternative was primarily due to *fastq_quality_filter* lacking the ability to run jobs in parallel.

Seqpac’s *make_trim* function generated very similar sequence counts compared to the *cutadapt/fastq_quality_filter* alternative (Figure 2B-C). We noticed, however, that *make_trim* generated slightly higher counts for some sequences (arrows in Figure 2B). Manually searching for these sequences across the original and trimmed fastq files showed that one explanation was that cutadapt failed to identify concatemer adaptor sequences. Concatemer (chimeric) adaptors are found in small quantity in most experiments, and are technical constructs where an incomplete adaptor associates with a complete adaptor during synthesis (34).

### 2.3 The low-level evidence filter

The *make_counts* function contains a low-level filtering module, here called an evidence filter. In default settings, it simply filters sequences that fails to replicate across two independent samples. Even in small experiments, such as the Kang et al. dataset, such filtering dramatically increases performance by reducing noise from extremely rare transcripts/degradation products (Figure 2C). Our experience is that such evidence filter often results in less than half the sequence diversity (number of unique read sequences; lower bars Figure 2C), while maintaining most of the sequencing depth (total number of reads; upper bars Figure 2C).

To illustrate this further, true sequence diversity—that can be replicated and is not due to technical bias—should expect to rise when sRNA from two species is mixed, which is also the case in the Kang et al. dataset (percentages in Figure 2C).

Nonetheless, the evidence filter in *make_counts* can both be disabled (e.g. in single sample/replicate experiments) or intensified (e.g. to increase performance in very large datasets). In addition, confirming our initial observation that seqpac’s *make_trim* function was better in identifying adaptor artifacts, such as concatemer adaptors, *make_trim* identified more replicable unique sequences passing the evidence filter than the popular *cutadapt/fastq_quality_filter* workflow (Figure 2D).

### 3.1 Annotating sequence with seqpac

Seqpac provides two ways to annotate a sequence in a PAC object. Firstly, the reannotation workflow (Figure 3: step 1-3) aligns the trimmed read sequences in the PAC against reference sequences, for example a reference genome, sRNA database, or sequences from another experiment, such as the results from a piwi pull-down. This is done using the reannotation family of functions: *map_reanno, import_reanno, add_reanno* and *simplify_reanno*. Seqpac also provides a ‘backdoor function’, *PAC_mapper*, that quickly calls the reannotation workflow for mapping the sequences in the PAC object (see Results 6.1, 7.2).

Secondly, after aligning a PAC object to a reference genome, the genomic coordinates of PAC sequences can be overlapped with coordinates of already annotated genomic features (Figure 3: step 4). This is done by the *PAC_gtf* function (Figure 3; step 4). Thus, by annotating using *PAC_gtf*, users can mimic a feature-based counting strategy, while saving the sequence integrity of the trimmed fastq-file in the PAC object.

### 3.2 Bowtie mapping using the *map_reanno* function

The reannotation workflow (Figure 3: step 1-3) depends on Bowtie (20) for sequence alignment, and therefore needs Bowtie indexes for the input fasta references. Similar to the adaptor trimming, *map_reanno* calls Bowtie either internally or externally, through the Rbowtie package (35) or a system call, respectively. The function can parse either seqpac standard or user provided options to Bowtie. It also calls a secondary function, *import_reanno*, which controls the import options from the Bowtie output files. Options involve for example whether coordinates and fasta sequence names should be reported, or only hit-or-no-hit. This is convenient for large repetitive sRNA references that may generate massive files if everything is reported (e.g. pirBase for humans and flies).

The *map_reanno* function runs multiple align/import cycles (Figure 3: step 1). After each cycle, imported data are saved as Rdata files, and only sequences without an alignment to any of the references will proceed to the next cycle. Each proceeding cycle allows for one additional mismatch until the user-defined max mismatches (or the Bowtie limit of 3 mismatches) has been reached. Reannotating only no-hit sequences in proceeding cycles not only guarantees that only the best hits are reported. Since system demands per sequence increases with each added mismatch, it also significantly increases performance as only the minimum number of sequences are aligned in each mismatch cycle. Importantly, if a sequence aligns to two references, both references will be reported for that cycle. Thus, unlike feature-based counting where such multimapping issues must be resolved already when reads are counted, users of seqpac can decide to discriminate between annotations at any stage in the analysis.

### 3.3 Annotating a PAC object using the *add_reanno* and *simplify_reanno*

Next, using the *add_reanno* function the Rdata files from each mismatch cycle is read into R and organized into a reanno object (Figure 3: step 2). For efficient access, this object is generated as a series of tibbles available in the tibble;tidyverse (36) package. Using a list of search terms, *add_reanno* consolidates the fasta sequence names into short character strings, which can be used as factors in downstream analysis. Search terms are constructed using regular expressions. A match will be reported as the reference name together with the search term. For example, if two references named mirbase and *ensembl_ncrna* were used as input for map_reanno, a search term list constructed as, list(mirbase=’mir’, ensembl=c(‘snoRNA’, ‘tRNÁ), will result in matches being returned as ‘mirbase:mir’, ‘ensembl:snoRNA’ and ‘ensembl:tRNA’. The user may choose if search terms must catch all reference hits, or if failure to match a search term should be returned as ‘other’ (e.g. ‘ensembl:other’).

Neither *map_reanno* nor *add_reanno* discriminates between references. Thus, if PAC sequences align to multiple references, all alignments and search matches will be reported (e.g. ‘mirbase:mir|ensembl:other’), but only if they align in the same mismatch cycle. For better transparency and reproducibility of sRNA experiments, we recommend that analysis is performed on a class-by-class basis as far as possible.

Nonetheless, hierarchical discrimination is often the only option to resolve some issues with pseudoreplication when multiple classes of sRNA are simultaneously analyzed. This is because the same sequence sometimes appears in multiple reference databases, and therefore obtains multiple classifications, such as both piRNA and miRNA. The purpose of the *simplify_reanno* function is therefore to hierarchically discriminate between search matches generated by the *add_reanno* function (Figure 3: step 3).

Importantly, since the seqpac workflow introduces simplified hierarchical classifications late in the annotation process, users can quickly set alternative hierarchies by just reapplying the *simplify_reanno* function. Unlike feature-based counting, the seqpac workflow therefore makes it easier to observe the effects of changes to the hierarchy. In addition, since seqpac maintains sequence integrity, users may at any time blast candidate sequences at their favorite genome browsers, to verify that the correct classification was made and to get additional information about the candidate.

### 3.4 Annotating genomic coordinates using PAC_gtf

When the reannotation workflow runs using the import=‘genome’ mode, the reference coordinates for each PAC sequence will be imported into the reanno object and later added to the PAC annotation table. These coordinates can be parsed to the *PAC_gtf* function as an alternative way to obtain PAC sequence annotations (Figure 3: step 4). This function uses gtf/gff formatted files that contains coordinates of genomic features and are available at many popular databases, such as Ensembl (37).

PAC_gtf simply overlaps PAC genomic coordinates with the gtf/gff coordinates using functions in the GenomicRanges package (38). It provides the user options on what information in the gtf to consolidate. Two predefined tracks, specifically expecting repeatMasker (39) and Ensembl (37) gtf files, are available besides a custom option.

### 3.5 Example: Reannotation workflow using the Kang et al. dataset

To exemplify seqpac’s reannotation workflow and plotting functions we ran multi-species mapping using the PAC object generated from the Kang et al. 2018 dataset (22) (presented in Figure 2). This involved parallel mapping to both the human (hg38) and fruit fly (dm6) genomes, as well as species specific versions of mirBase (miRNA) (40), pirBase (piRNA) (41), GtRNAdb (tRNA) (42) and ensembl (many types of ncRNA) databases (37). The hierarchy was set to rRNA > tRNA > miRNA > snoRNA > snRNA > lncRNA > piRNA, indicating that rRNA was most prioritized and piRNA was least prioritized. Mapping was carried out allowing for up to 3 mismatches.

As expected, the test clearly discriminated between human and fly samples in terms of genome alignment, and correctly accounted for the expected genomic ratios when samples from these two species had been mixed (Figure 4A). The human proportion of the dataset was more affected by perfect matching, which is expected due to more outbreeding in the population, but both species gain almost 100% ‘mappability’ when mismatches were allowed. The fruit fly proportion of the dataset was strongly enriched with an rRNA sized to 30 nt (Figure 4B). Blasting this sequence showed that it was identical to the complete 2S rRNA subunit. This was expected since Kang et al. did not to report of any method that depletes 2S rRNA prior to library construction, which is commonly done in fruit fly experiments (43, 44). The human proportion of the dataset was instead enriched with miRNA with the expected size of 22 nt (Figure 4B). There was also a T bias in the expected range between 22-25 that may indicate piRNA (Figure 4C). Nonetheless, the proportion of piRNA classification was lower than the T bias (Figure 4B/C), which suggests that some piRNA may have been classified as miRNA given that miRNA was prioritized in the hierarchy.

### 4.1 Subsetting and grouping data using targeting objects

Seqpac applies an innovative strategy for extracting sample groups and sequence classifications for filtering, plotting and statistical purposes. This involves small targeting objects constructed as a list with two inputs (Figure 5). The first being a character string naming a target column in a specific table held by the PAC object, while the other is a character vector naming the target entries of the target column. Importantly, the name of the targeting object itself pinpoints to which PAC table that should be targeted. Thus, if a function has a ‘pheno_target=’ input, a targeting object naming a column in the phenotype table can be used to subdivide the data. Similarly, if an ‘anno_target=’ input option is available then columns in the annotation table can be targeted. The second entry of a targeting object is often order sensitive. Thus, if users want the sample groups to appear in a specific order in a graph, they only need to provide that order in the second entry of the *pheno_target* object (Figure 5).

**Figure 5.**
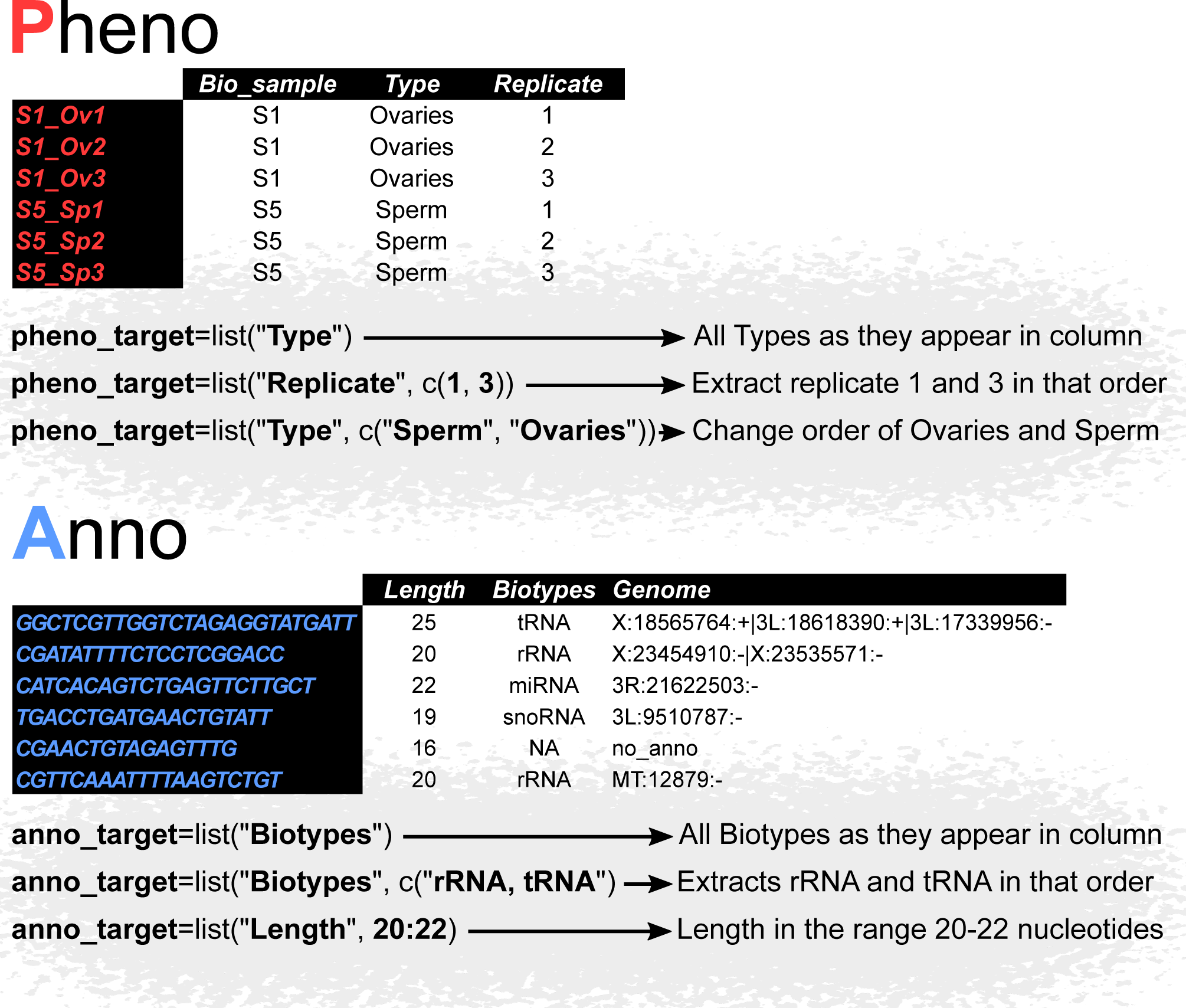
The principles for working with target objects. Many seqpac functions applies a novel system for grouping and sub-dividing samples (Pheno) and sequences (Anno) in a PAC object. This system relies on small target objects, which targets information either in the Pheno (pheno_target) or Anno (anno_target) tables. A target object is a list with two-character inputs. The first pointing to a column in the target table, and the second to the entries of that column.

As an example, when using the *PAC_pie* to generate the pie charts in Figure 4A, we used an *anno_target* for a column in the Anno table containing the four different genome classifications (second entry order: “No alignment”, “Fly”, “Human” and “Both fly and human”). Similarly, when generating the size distribution histograms in Figure 4B, we used an *anno_target* for a column in Anno holding the sRNA classifications generated by the *simplify_reanno* function.

In a few cases, seqpac functions use targeting objects for other seqpac objects, such as a PAC summary table (see Results 5.3). While the principle of these objects is similar to the *pheno_target* and *anno_target* objects, they may have differences that are carefully described in the manual to each function.

### 5.1 Overview preprocessing, summarization and statistical analysis

With or without advanced annotations, PAC objects can be filtered (PAC_filter, PAC_filtsep), normalized (PAC_norm), and summarized (PAC_summarize) using seqpac internal functions. More advanced statistical wrappers immediately compatible with PAC objects are also available (e.g. PAC_deseq, PAC_pca).

### 5.2 Filtering

PAC_filter and *PAC_filtersep* handles filtering and subsetting of PAC objects. With PAC_filter, users can subset the PAC object by targeting columns in the Pheno and Anno tables using the *pheno_target* and *anno_target* options (see Results 4.1). A filter that extracts sequences that have reached a percent coverage over a certain threshold is also available for both raw and normalized counts. This can for example be used for the popular ‘20 counts in 50% of samples’ filter. *PAC_filter* can also plot a graph that shows the impact on the data at different thresholds. Conveniently, seqpac provides a separate function PAC_filtersep, that extracts sequences reaching a coverage threshold within sample groups. The output can directly be used to construct Wenn-diagrams, for example visualizing the sequence overlap that reach 100 cpm within two sample groups. It can also be applied for more advanced filters, like removing read sequences that do not reach 20 counts across all samples within a group.

### 5.3 Normalize, summarize and statistical analysis

While the standard structure of a PAC list object contains three tables—Pheno, Anno and Counts—it may hold any number of objects as long as they do not have the same names as the standard objects, just like a regular list. There are, however, two more standard objects that are added to the PAC object later in the analysis: the norm list containing normalized counts tables, and the summary list that contains summarized tables (Figure 6). It is easy to visualize these objects as two separate ‘folders’ within a PAC object.

**Figure 6.**
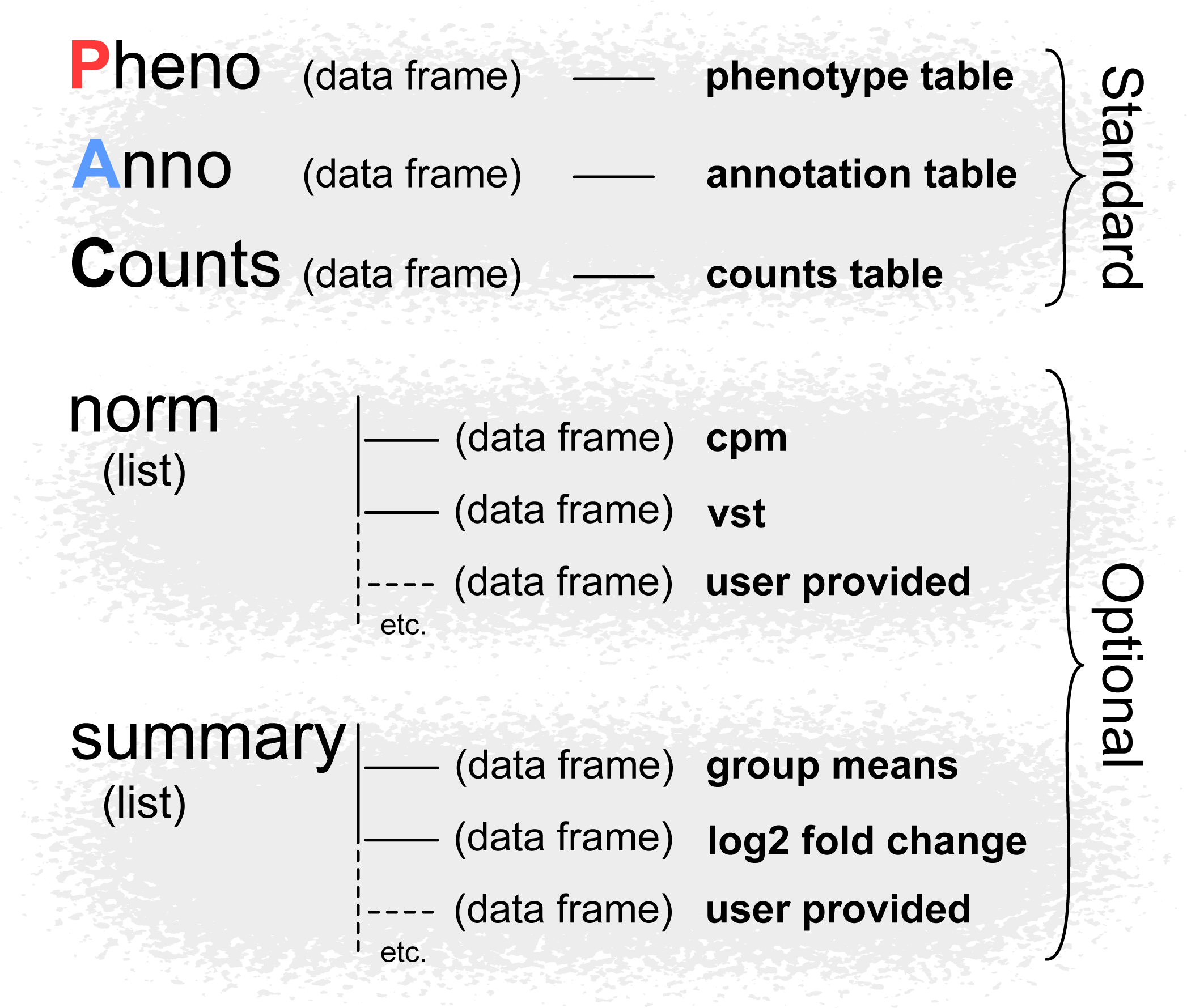
Advanced PAC objects. All PAC objects must contain Pheno, Anno and Counts tables. Many seqpac functions may optionally use data stored in two additional PAC objects: the norm and summary lists. The norm contains tables of normalized counts having identical row and column names as Counts. The summary contains tables with identical row names as Counts, but column names based on aggregates over the Counts columns (e.g. group means). These PAC ’folders’ can be generated by the *PAC_norm* or *PAC_summary* functions, but users are encouraged to provide their own tables.

**Figure 7.**
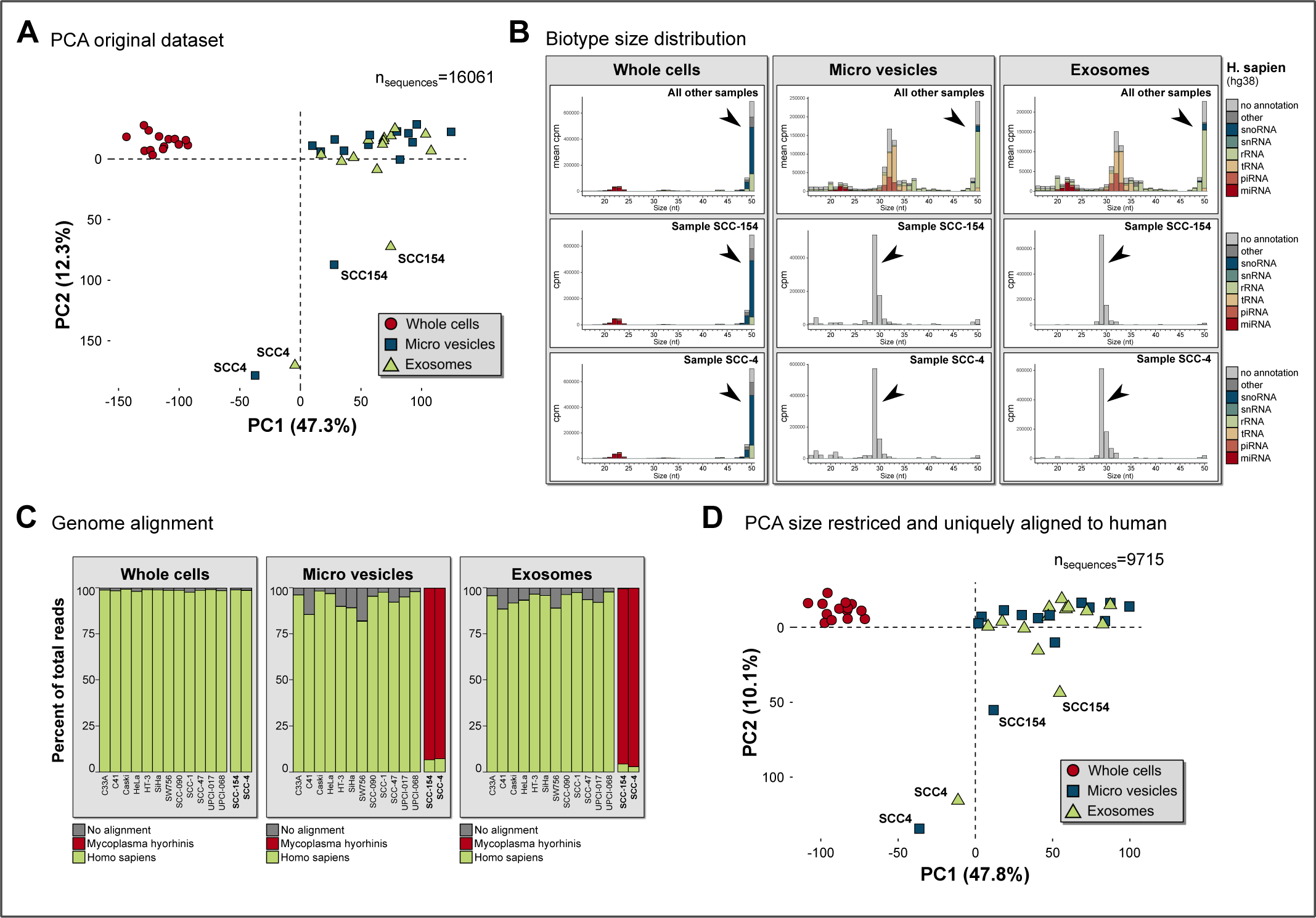
Seqpac example 1: Identifying contaminants by sequence-based counting. We generated a PAC object from a public dataset (SRA access: PRJNA666144). This study used a feature-based counting strategy to examine the sRNA in cells, and their extracellular vesicles, of 14 cervical and head/neck cancer cell lines. (A) Scatter plot generated by the *PAC_pca* function after vst normalization using *PAC_norm*. The two first principle components identify extracellular vesicles from SCC4 and SCC154 cells as outliers. (B) Size distribution histograms generated by the *PAC_sizedist* function. Most samples show high content of sRNA ≥ 50 nt, except for the SCC4 and SCC154 outliers, which are enriched with 29 nt fragments with no annotation in humans. (C) Bar graphs generated by the *PAC_stackbar* function after reannotating the PAC object against the human and Mycoplasma hyorhinis (contaminant) genomes show high content of Mycoplasma in outliers. (D) Scatter plot generated by *PAC_pca* after removing all non-human sequences with the *PAC_filter* function, only including sRNA between 16-45 nt, and re-normalizing the data with *PAC_norm*. No big differences from the original dataset (A).

**Figure 8.**
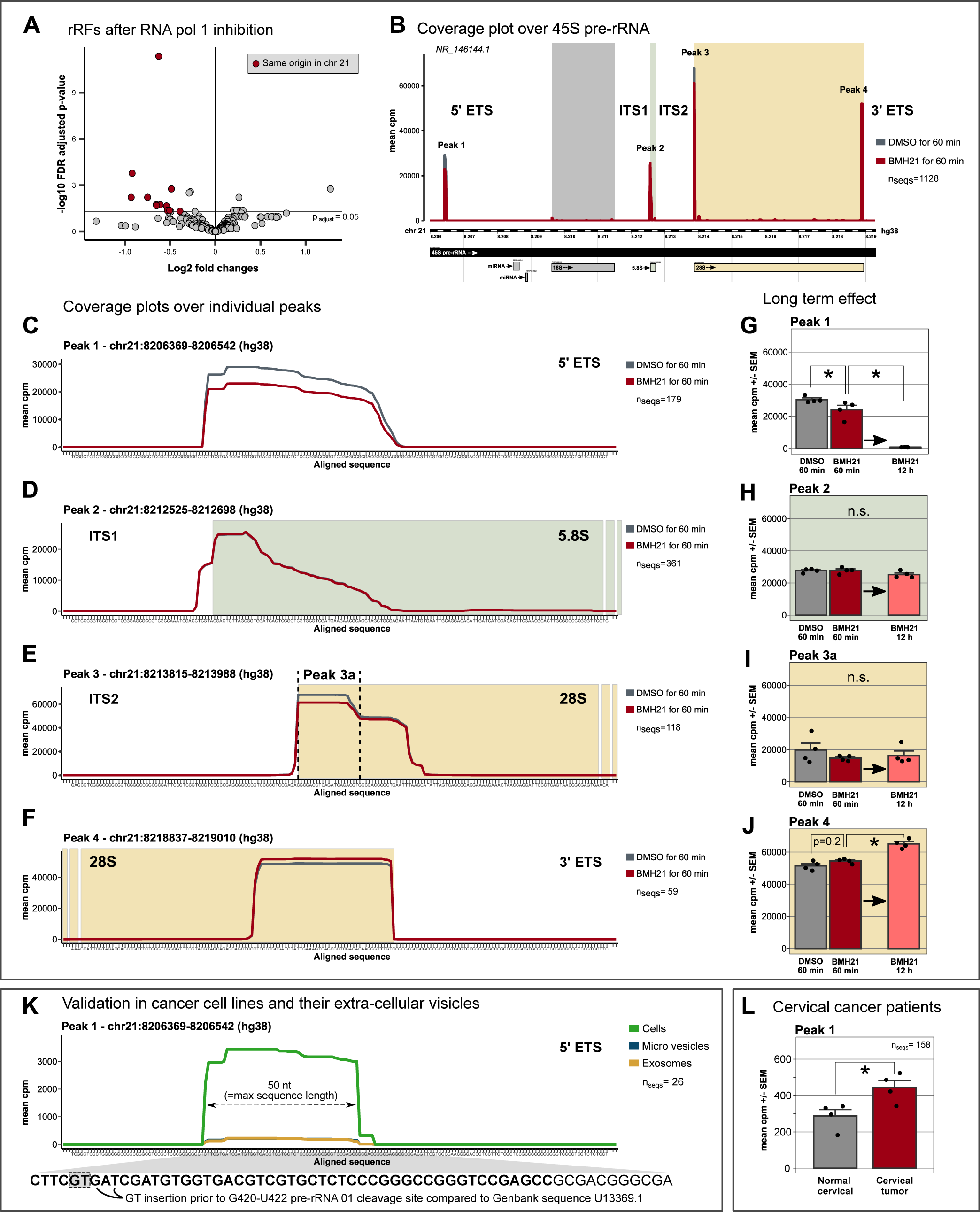
Seqpac’s sRNA coverage functions reveal novel rRNA derived fragment (rRF) in cancer. RNA polymerase I (Pol I) transcribes the 47/45S pre-rRNA and has been targeted in anti-cancer treatment. Small RNA analysis of HeLa cells either exposed to an RNA polymerase I inhibitor (BMH21) or DMSO (control) for 60 min (A-J)(SRA access: PRJNA708219). Volcano plot (A) from a differential expression analysis using the *PAC_deseq* function showing down-regulated rRFs. Red indicates related sequences that primarily originates from an rRNA cluster on chr 21. Coverage plot (B) using the *PAC_mapper* and *PAC_covplot* functions of an 45S pre-rRNA on chr 21 (GenBank: NR_146144.1). Shows 4 major peaks affected differently by BMH21. Zoomed in coverage plots of Peak 1-4 (C-F). Peak 1 (C) shows a novel 5’ ETS rRF downregulated by treatment. Peak 2 (D) shows a plausibly degraded fragment. Peak 3 (E) contains two overlapping rRFs where only the small (Peak 3a) varies by treatment. Peak 4 (F) shows a rRF possibly upregulated by treatment. Bar graphs (G-I) showing that 12h exposure to BMH21 amplifies the effects in Peak 1 and Peak 4, but not in Peak 2 and Peak 3. Coverage plots (K) validating the 5’ ETS rRF of Peak 1 in cancer cells, and to a lesser degree in extra-cellular vesicles, using the Tong et al. dataset from Figure 7 (without the contaminated SCC4 and SCC154 cell lines). The zoomed in genomic sequence (bottom) shows the main fragment from (C) where the longest fragment from Tong et al. is presented as bold letters. Dotted grey box indicate GT-insertion compared to a commonly studied 47S pre-rRNA (GenBank: U13369.1). Bar graphs (L) showing an up-regulation of 5’ ETS rRF related fragments in cervical tumor samples published by Xu et al. (SRA access: PRJNA607023). Number of summed sequences that were found in the target regions are indicated by nseqs. Mann-Whitney U tests indicated by * p<0.05 and ^#^ p<0.1 significance levels. SEM = Standard error of the mean.

PAC_norm provides a few common normalization methods, like the simple reads/counts per million that standardize each sample against their total counts. It currently also maintains a wrapper for the rlog and vst functions of the DESeq2 package (29), that automatically will prepare the PAC counts table for a transformation blinded against experimental groups. Users are, however, encouraged to provide their own normalization tables. As long as the table contains the same sequence (row) and sample (column) names as the Counts table, and are stored in the norm list (‘folder’) of the PAC object, seqpac functions with a norm input option will automatically search the norm folder for a matching name.

PAC_summary generates simple group summaries, like means, standard deviations, standard errors, percent group differences and log2 fold changes. It can be applied to both raw counts, as well as normalized counts by naming a table in the PAC norm list/folder using the norm input option. The grouping of samples is controlled by a *pheno_target* object. *PAC_summary* does not maintain an *anno_target* option since summaries over annotations would result in loss of sequence integrity (= feature-based counts). Summarizing data across both phenotype and annotation is instead handled by individual functions, or by subdividing the whole PAC file using the *PAC_filter* function prior to running PAC_summary.

For more advanced statistical analysis seqpac provides a convenient function, PAC_deseq, that allow users to import a PAC object into DESeq2 (29). This function automatically generates a report containing organized top tables, volcano-plots and p-value distribution histograms. Further, seqpac contains the *PAC_pca* function that performs a principle component analysis (PCA) with aid of the FactoMineR and factoextra packages (45, 46). This function returns scatter plots of the main components annotated using either a *pheno_target* or anno_target. Lastly, *PAC_saturation* performs and plots the results of a sequence saturation analysis.

This is often used for checking that satisfactory sequencing depths have been reached, where few new sequences are predicted given a hypothetical increase in the sequencing depth.

### 6.1 Advanced classification and visualization

In the quick reference presented in Table 1 a selection of visualization functions is briefly presented. In common for most of them are the option to use *pheno_target* and/or *anno_target* objects for grouping and ordering different plots (as described in Figure 5). Seqpac plots are primarily generated using the ggplot2 package (47) and outputs are often saved as lists with summarized data and graphs. As with the other seqpac functions, outputs are described in detail in the functions’ manuals.

Since seqpac’s reannotation workflow provides a powerful and quick pipeline for sequence annotation, we have also included a ‘back-door’ function, PAC_mapper. This function is ideal for detailed mapping of smaller fasta references, such as a list of tRNAs, the 45S pre-rRNA subunit, the mitochondrial genome, or simply a specific genomic region download as a fasta from a genome browser. Conveniently, if a Bowtie index is missing for a fasta reference, *PAC_mapper* will automatically generate that index, making the alignment of a new fasta reference highly efficient. The output of *PAC_mapper* is a map object, which is simply a list where each entry refers to a specific sequence in the fasta reference and where the coordinates of all PAC sequences that mapped the reference sequence is reported (e.g. the mapping coordinates of PAC sequences aligning to a specific tRNA). This map object along with the original PAC object can then be fed to the *PAC_covplot* function to generate coverage plots across the fasta reference, as exemplified in Figure 8. As we have illustrated before, such coverage plots are well suited for characterizing tRNA and rRNA fragmentation (8,44,48), as well as mitochondrial RNA (48).

Lastly, using the map object the *map_rangetype* function can generate more advanced classifications such as 5’, i’ and 3’ tRFs or tRNA halves, previously best demonstrated in MINTbase and MINTmap (11). Nonetheless, the MINTmap/MINTbase suite is only readily available for human tRNA classification. Seqpac’s *PAC_mapper* and *map_rangetype* functions fills this gap and expands the possibility for discovering novel tRNA fragment classes in any species. With the *map_rangetype* function it is easy to classify sequences in the map object in relation to where the alignment starts or ends in the reference sequence. This is done by either defining different ranges (e.g. classifying a fragment as 5’ if it starts within the first 3 nt of a tRNA), or a percentage zone (e.g. classifying a fragment as a half if it ends or starts within 45-55% of a tRNA). Even better, *map_rangetype* may use ss files, which is a format commonly used for storing information about secondary structures such as tRNA loops. Thus, using this option, users can classify fragments in relation to for example cleavage within a specific loop. We used this strategy to identify a diet-sensitive tRNA derived fragment in human sperm, that we called nuclear internal T-loop tRNA derived RNA (nitRNA)(48). The *PAC_trna* function plots range-classified tRNAs mimicking some graphs presented in that paper.

### 7.1 Seqpac example 1: Identifying contaminants by sequence-based counting

To further illustrate the strengths of seqpac, we reanalyzed a recently published dataset by Tong et al. 2020 (25) (SRA access: SRP285629). This dataset contains 42 fastq files originating from 14 human cancer cell lines, where RNA was extracted from cells, as well as exosomes and microvesicles of these cells. Extra-cellular vesicles—such as microvesicles and exosomes—are cellular excretion particles produced by cells’ plasma membrane. They are found from a variety of cells— including tumor cells—in peripheral body fluids (49). Therefore, characterizing the sRNA content of extracellular vesicles from cancer cell lines may reveal novel diagnostic/prognostic biomarkers.

We generated a PAC object from this dataset. The sequencing was done on an Illumina HiSeq3000 sequencer with a flow cell kit generating read lengths of only 50 nt. From our experience, we do not recommend generating sRNA-seq data with read lengths shorter than 75 nt. Longer reads allow for inter-adaptor length validation, where detecting the opposite adaptor sequence in the read guarantees that it originated from short RNA and not from long RNA. Thus, unless controlling for sequence length in downstream analysis, sRNA experiments with very short reads may be severely influenced by long RNA. To investigate if this was a problem in the Tong et al. study, we therefore included all read lengths in the analysis.

Tong et al. (25) used a feature-based counting strategy. This strategy first aligns sequences to a reference genome, often allowing for multiple mismatches, and discards sequences that fails to align. Counts are then based on the overlaps between the genomic coordinates of the reads and the genomic coordinates of known sRNA. This poses several problems. The nucleotide sequence of some sRNA may be post-transcriptionally modified, such as 3’ fragments of mature tRNAs. These may be discarded since they fail to align with the reference genome. Further, allowing for mismatches without knowing where those mismatches occur and pool related sequences with and without mismatch alignments into the same feature, can hide information about sRNA subtypes and remove traces of post-transcriptional modifications hidden within the reverse-transcription signature (50–52). Many sRNAs are also highly repetitive, such as most rRNA derived fragments, and may thereby map to multiple genomic regions. In feature-based counting strategies this is often solved by randomly assigning such reads to one of the multimapping regions.

Together, this completely breaks sequence integrity making it difficult to interpret the results.

Some of these issues with feature-based counting can be illustrated with the Tong et al. dataset (25) using seqpac’s workflow. It must be emphasized, however, that our critic is not specifically aimed against Tong et al., whose work we admire, but rather against the feature-based counting strategies that hundreds of studies have been using.

By applying a PCA we confirmed what the original authors reported that cells were very different from extra-cellular vesicles (Figure 7A). We also observed that the extra-cellular vesicles from two specific cell lines—SCC4 and SCC154—were different to the other samples. Size distribution histograms immediately identified two problems (Figure 7B). Firstly, most read sequences were ≥ 50 nt. Since Tong et al. reported that the majority of sRNA from cells came from snoRNA, and sRNA from extra-cellular vesicles came from rRNA, it indicates that their analysis did not account for sequence length. This is because most snoRNA and rRNA are found in the >= 50 nt segment. Thus, the sRNA class proportions that was reported may involve long RNA, possibly including full-length rRNA and tRNA.

Secondly, the extra-cellular vesicles from SCC4 and SCC154 failed almost completely to align with known human sRNAs (Figure 7B). Since seqpac maintains sequence integrity, we blasted a small selection of these non-annotating sequences at NCBI (53). The result strongly indicated that most reads originated from the Mycoplasma hyorhinis genome. Since this is a common contaminant in cell cultures (54), we ran this Mycoplasma genome in parallel to the human genome in the seqpac’s reannotaion workflow, thereby picking the best possible alignment from either of them. This showed that all vesicle samples from SCC4 and SCC154—the same samples that explained one of the main components in the PCA—suffered severely from Mycoplasma contamination (Figure 7C).

Now, critics may argue that a feature-based counting strategy should have corrected for this contamination, since reads that fail to align against the human genome will automatically be removed prior to counting. Thus, Mycoplasmic reads should not have affected the results, since normalization of the counts was made after their removal.

We tested this assumption using seqpac functions. With the output from the two-genome reannotation workflow, we used the *PAC_filter* function to remove all sequences that mapped to the Mycoplasma genome, and only kept reads that mapped to the human genome. Then we re-normalized the dataset using the *PAC_norm* function and made a new PCA. Removing nearly 6500 sequences, and keeping only sequences exclusive to the human genome, had very limited effects on the results (Figure 7D). This strongly indicates that the effect of the contamination remained even after removing the contaminating sequences. Importantly, this bias may have gone unnoticed if we would have used a feature-based counting strategy, since contaminating sequences would have been removed prior to counting.

Together, this illustrates how seqpac quickly provides panoptic views of data integrity, which is essential for analytical transparency and correct downstream interpretations.

### 7.2 Seqpac example 2: Novel rRNA-derived sRNA affected by anticancer treatment

In cancer research, non-coding RNA has been studied not only for diagnostic and prognostic purposes, but also for therapeutic purposes (55). Of particular interest, rRNA synthesis is commonly exaggerated in tumor cells (56). Synthesis involves transcription of 47S/45S pre-rRNA genes by RNA polymerase I at specific repetitive clusters in the genome (57). Over a series of precursors, pre-rRNA is turned into the active mature rRNA subunits 28S, 18S and 5.8S (58, 59). Inhibiting RNA polymerase I (RNA pol I) has been proposed as a possible anticancer treatment, where one of the most promising candidates have been the BMH21 compound (60). However, little is known about sRNA generated from the pre-rRNA and their potential role in cancer.

We, therefore, used seqpac to detect novel sRNA originating from pre-rRNA, hypothesizing that inhibiting RNA pol I would result in fewer rRFs. For this we conducted a small experiment by exposing HeLa cells—which originates from cervical cancer cells—to BMH21. The exposure time was set to 60 min, and as control we used DMSO. RNA from purified cells was then prepared for sRNA-seq, which resulted in fastq files with 75 nt reads. From this raw data we generated an annotated and filtered PAC object using only seqpac functions.

Using the *PAC_deseq* (see Results 5.3) function, we performed a differential expression analysis only including highly expressed sRNA mapping to rRNA reference sequences. This showed that only 60 min of BMH21 exposure was enough to affect rRNA fragmentation (Figure 8A). Perhaps unexpectedly, not all were downregulated by inhibiting RNA polymerase I. In fact, closer examination revealed that most down-regulated sequences were related (Figure 8A; Supplementary table S1), suggesting a single origin within an rRNA cluster on chromosome 21. We, therefore, used the *PAC_mapper* and *PAC_covplot* functions (see Results 6.1) to visualize the impact of BMH21 over a pre-rRNA 45S gene on chromosome 21 (GenBank: NR_146144.1). This revealed 4 major rRFs (Peak 1, 2, 3, 4 in Figure 8B), where the related fragments from Figure 8A all aligned to Peak 1. For more detailed analysis, we downloaded the sequences of the DNA immediately neighboring these peaks from the UCSC genome browser and ran the sequences as a fasta reference file in the *PAC_mapper* and *PAC_covplot* functions (Supplementary file S2). This revealed what appeared to be a single large down-regulated fragment in Peak 1 (Figure 8C), an unaffected possibly degraded fragment in Peak 2 (Figure 8D), two separate fragments in Peak 3 where only the shorter and less expressed fragment might have been affected by BMH21 (Peak 3a in Figure 8E), and one single fragment in Peak 4 that seemed slightly up-regulated following BMH21 treatment.

To better understand the relevance of these changes we summed the cpm of all fragments mapping to each peak and performed a non-parametric Mann-Whitney U test. For this analysis we also included a third group of samples that had been exposed to BMH21 for 12 hours, to explore if any of the effects of BMH21 were amplified following long-term exposure. Astonishingly, after 12 h exposure, fragments of Peak 1 had almost completely disappeared (Figure 8F). This was not due to an experimental failure since Peak 2 and Peak 3a were unaffected by the long-term treatment (Figure 8G-H). In fact, Peak 4 fragments even showed a significant up-regulation (Figure 8I). Thus, the effects observed in Peak 1 and Peak 4 were amplified by long-term exposure, but in two opposite directions.

For the Peak 2 and Peak 3 rRNA fragments we have previously observed similar fragments in human sperm (8), and similar fragments located to the 5’ ends of the 5.8S and 28S subunits in fruit fly embryos (44). This is also true for the 3’ fragments of the 28S subunit (Peak 4), even though we never have observed such expression levels as we see in the HeLa cells. To our knowledge, however, highly expressed sRNA fragments from the Peak 1 region—in the 5’ external transcribed spacers (ETS)—have never been described. To understand the 5’ ETS rRF better, we performed a multi-species blast of the main sequence at NCBI to identify similar GenBank entries. This showed many alignments to ribosomal precursors in humans, one identical sequence in the Chimpanzee, and a few similar sequences in the Gorilla (Supplementary Figure S1). Thus, this 5’ ETS rRF has only evolved in our closest relatives.

Confident that the 5’ ETS rRF was a human sRNA, we searched for this fragment in the Tong et al. 2020 dataset. Despite only having read lengths of 50 nt to our disposal (see Results 7.1), where 5’ ETS rRF of Peak 1 was 61 nt, we found clear traces of this rRF (Figure 8J). Furthermore, to explore the clinical relevance of this finding we downloaded the Xu et al. dataset (26, 27). Here sRNA was extract from confirmed cervical tumors and samples from normal cervix. Results indicated that the 5’ ETS rRF was upregulated in cancer patients (Figure 8K). Together this suggests that our novel rRF—validated by the seqpac workflow in multiple unrelated datasets—may be targeted for diagnostic and prognostic purposes during cancer treatment.

## DISCUSSION

Here we presented a novel and innovative bioinformatic tool—seqpac—that makes advanced sRNA analysis from genome-scale sequencing data more accessible and transparent. The workflow is completely integrated with R, from trimming the adaptor sequences to generating plots. We showed that seqpac’s trimming function performs as well as, or even better, than trimming using standard tools outside R. We further presented the PAC object, which builds a framework of phenotypic information (P) and sequence annotations (A) around a table based on sequence counts (C). Using published data we showed that a sequence-based counting strategy—in contrast to feature-based counting that is more commonly used—diminishes the risk of mistakes in downstream analysis. We demonstrated the strength of maintaining sequence integrity to enable re-annotation of sequences across species and classes of sRNA at any point in the analysis. Lastly, we showed how seqpac can be used for sRNA discovery in cancer research by the discovery of a novel rRNA derived fragment (rRF) that were down-regulated by anti-cancer treatment in vitro and up-regulated in tumors of cervical cancer patients.

Seqpac is available at github (https://github.com/Danis102/seqpac). As the whole workflow, from adaptor trimming to mapping and plotting, are integrated in R it runs on common computer platforms, including Windows, Mac and Linux. It comes with a complete collection of function manuals and a vignette that guides the user in how to apply the default workflow using a fastq test dataset that are included with the package. R scripts that we used to generate many of the results presented in this paper are available in Supplementary file S1.

It must be emphasized that seqpac is primarily designed for sRNA sequence analysis. This means that it does not currently supports paired-end sequencing, which is commonly applied for long RNA sequencing. Paired-end sequencing is not required for most sRNA applications where the target sequence lengths seldom exceed 75 nt. As we have demonstrated in this paper, too short reads—as those generated using the 50-cycle flow cell kits available for MiSeq, NextSeq1000 and HiSeq2500/3000/4000—should be avoided. Without some excessive sequence in which the 3’ adaptor can be detected, it is difficult to reliably discriminate medium length sRNA (such our novel 5’ ETS rRF) from unintentionally included longer RNA.

We see, however, many advantages to use sequence-based counting also in long RNA sequence analysis, for example to easily extract sequences annotating to a candidate mRNA and check for possible genetic variants. Coverage plots, similar to what we describe for the 45S pre-rRNA (Figure 8B) would also be applicable for mRNA coverage to visualize splice variants and intronic transcription. Even though we hope to develop long RNA analysis in future updates of seqpac, there are currently a few technical constraints that needs to be resolved.

As mentioned, paired-end reads are not supported either for trimming or counting. In addition, while Bowtie (20) is still the most popular aligner for sRNA, it does not support indel mapping. While this is not a great problem if sequence integrity is maintained and candidate sequences subsequently can be blasted to detect any slip through, this problem are slightly more announced in samples differing much from their reference genomes, such as cancer cell-lines. A likely reason for Bowtie’s popularity in sRNA community is because it is reliable with short sequence alignments. For instance, we initially tried to integrate the Rsubreads package (61) in seqpac’s workflow, which applies a highly efficient ‘seed-and-vote’ mapping algorithm. However, for certain read lengths we consistently experienced failure to correctly vote for the best alignment, possibly as a consequence that too few seeds were covering the read. We will off-course explore more efficient alternatives to Bowtie in the future.

By using the sequence-based approach of seqpac, we have discovered a novel rRNA derived sRNA (rRF) in the 5’ ETS of 45S pre-rRNA. This rRF responds negatively to anticancer treatment and are up-regulated in tumors. The scope of our study was not to dwell deep into the mechanism and clinical potential of this rRF. To our knowledge, however, this fragment has not been described before, and from our experience sRNA in the 5’ ETS of pre-rRNAs are rare. This, together with the insight that the sequence is relatively unique to humans (with only some homology in Chimpanzees and Gorillas), makes it a good target for future studies on biomarkers in cancer treatment and diagnosis. In our HeLa cell experiment, the main fragment was 61 nt, which indicates a unique fragment given that we had a maximum read length of 75 nt. Even though the methods used in Tong et al. (25) and Xu et al. (26, 27) were restricted to a maximum read length of 50 nt, we found traces of this fragment in the pile of fragments with unverifiable length of ≥ 50 nt. It must be emphasized, however, that we tried to validate the 5’ ETS rRF in yet another dataset, Snoek et al. (62) (SRA accession: PRJNA413777), but here we failed to detect anything in the 5’ EST region. The Snoek et al. dataset is so far the largest public sRNA dataset from cervical cancer patients. In this study, samples were collected by participants themselves, which may explain much higher rRF variability and excessive number of short fragments (< 20 nt), compared to the other datasets (Supplementary file S1). Importantly, in contrast to the other datasets that used the NEBNext Small RNA Library Prep kit, Snoek et al. used the Illumina TruSeq Small RNA Library Preparation Kit. We and others have consistently shown that these two popular kits perform differently with regard to sRNA coverage (48,63–65). Thus, future studies targeting this rRF must consider the choice of chemistry, and maybe even apply advanced protocols for better coverage (44, 66).

The 5’ ETS rRF should be located somewhere close to the 01 cleavage sites in the 5’ EST of 47S pre-rRNA. When aligning the 61 nt main fragment against the GenBank entry U13369.1—which have been commonly used to map Human pre-rRNA cleavage sites (59)—the 5’ ETS rRF only partly align. The coverage over this area in the U13369.1 pre-rRNA is also far from what we observed for the NR_146144.1 pre-rRNA, which was the GenBank sequence that we used for the chromosome 21 alignment in (Figure 8B; Supplementary File S2). Aligning the U13369.1 with NR_146144.1 reveals a “G-T” insertion in NR_146144.1, right between the C414-C416 and G420-U422 01 cleavage sites (67) (Figure 8K; Supplementary Figure S2). Since the 5’ ETS rRF contains this insertion, it strongly suggests that pre-rRNA cleavage has been affected at this locus. Therefore, beside investigating possible clinical values of this 5’ ETS rRF, future research may target mechanisms for how natural rRNA variants may give rise to novel sRNA, suggestively by investigating the interactions between post-transcriptional modifications, snoRNA and proteins at this locus.

In conclusion, the revolution of genome scale sequencing has not only brought enormous potential for unraveling life’s mysteries in health and disease. It has also created a gap between biology and technology. Consequently, research groups with primary biological or medical interests are often forced to rely on specialized programmers with limited understanding of the biology to handle their precious data. R has long been a platform where bridges between biology, statistics and programming are built. We have showed that building a transparent workflow in R for sRNA analysis, with the intention of making choices early in the analysis perspicuous, not only helps in detecting severe biases that would have otherwise gone undiscovered. It also provides the flexibility and panoptic view needed for advanced biological interpretations.

## Notes

https://github.com/Danis102/seqpac

https://www.ncbi.nlm.nih.gov/bioproject/PRJNA708219/

### Competing Interest Statement

The authors have declared no competing interest.

