## Supplementary Figures and Tables for "Seqpac: A New Framework for small RNA analysis in R using Sequence-Based Counts"

### Supplementary Table S1. Results differential expression analysis between BMH21 60 min and vs DMSO 60 min control

| *Sequence* | *size* | *base mean* | *log2 fold change* | *p-val (adjusted)* [A] | *hg38 alignment* [B] |
|---|---|---|---|---|---|
| **TTCGTGATCGATGTGGTGACGTCGTGCTCTCCCGGGCCGGGTCCGAGC** | 48 | 530.86 | -0.63 | 4.43E-12 | perfect |
| **TTCGTGATCGATGTGGTGACGTCGTGCTCTCCCGGGCCGGGTCCGAGCCGCGACGGGCGAGGG** | 63 | 1175.41 | -0.92 | 0.00017 | perfect |
| **ATCGATGTGGTGACGTCGTGCTCT** | 24 | 1278.9 | -0.49 | 0.0017 | perfect |
| GGCGCGGGACATGTGGCGTACGGAAG | 26 | 522.5 | 1.27 | 0.0017 | perfect |
| CGACTCTTAGCGGTGGATCACTCGGCTCGTGCGTCGATGAAGAACG | 46 | 960.93 | -0.28 | 0.0027 | perfect |
| CGCGACCTCAGATCAGACGTGGCGACCCGCTG | 32 | 2043.21 | -0.29 | 0.0031 | perfect |
| **TTCGTGATCGATGTGGTGACGTCGTGCTCTCCCGGGCCGGGTCCGAGCCGCGACGGGCGAGG** | 62 | 2591.06 | -0.93 | 0.0061 | perfect |
| **CTTCGTGATCGATGTGGTGACGTCGTGCTCTCCCGGGCCGGGTCCGAGCCGCGACGGGCGAGGG** | 64 | 593.59 | -0.75 | 0.0061 | perfect |
| GGGAGACCGCCTGGGAATACCGGGTGCTGTAGGCTTT | 37 | 1396.97 | 0.26 | 0.0061 | perfect |
| **CTTCGTGATCGATGTGGTGACGTCGTGCTCTCCCGGGCCG** | 40 | 465.9 | -0.61 | 0.019 | perfect |
| **TTCGTGATCGATGTGGTGACGTCGTGCTCTCCCGGGCCGGGTCCGAGCCGCGACGGGCGAG** | 61 | 3281.02 | -0.65 | 0.019 | perfect |
| **CTTCGTGATCGATGTGGTGACGTCGTGCTCTCCCGGGCCGGGTCCGAGCCGCGACGGGCGAG** | 62 | 5015.92 | -0.65 | 0.021 | perfect |
| **ATCGATGTGGTGACGTCGTGCTCTCCCGGGCCGGGTCCGAGCCGCGACGGGCG** | 53 | 432.09 | -0.54 | 0.022 | perfect |
| **CTTCGTGATCGATGTGGTGACGTCGTGCTCTCCCGGGCCGGGTCCGAGCCGCGACGGGCGA** | 61 | 6240.52 | -0.51 | 0.043 | perfect |
| CGTACGACTCTTAGCGGT | 18 | 784.86 | 0.27 | 0.044 | perfect |
| **CTTCGTGATCGATGTGGTGACGTCGTGCTCTCCCGGGCCGGGTCCGAGCCGCGACGGGCG** | 60 | 3931.47 | -0.53 | 0.044 | perfect |
| GATGGGAGACCGCCTGGGAATACCGGGTGCTGTAGGCTT | 39 | 573.74 | 0.31 | 0.044 | perfect |
| CTCGCTGCGGTCTATTGAAAGTCAGCCCTCGACACAAGGGTTTGT | 45 | 494.04 | 0.2 | 0.044 | 1 mismatch |
| GACTCTTAGCGGTGGATCACTCGGCTCGTGCGTCGA | 36 | 476.28 | 0.24 | 0.044 | perfect |
| CGCGACCTCAGATCAGACGTGGCGACCCGCTGAATTTAAGCAT | 43 | 774.49 | -0.5 | 0.05 | perfect |
| **TTCGTGATCGATGTGGTGACGTCGTGCTCTCCCGGGCCGGGTCCGAGCCGCGACGGGC** | 58 | 2307.88 | -0.39 | 0.05 | perfect |

Analysis was done using the PAC_deseq function after extracting read sequences aligning to rRNA reference sequences and obtaining at least 100 cpm in all sample within either BMH21 (treated) or DMSO (control) exposed for 60 min. In total 211 rRFs were analyzed. **Bold** indicates the related sequences (red dots) from Fig 8A.

[A] P-value corrected for false discovery rate by the Benjamini and Hochberg method.

[B] Genome alignments were done using the reannotation workflow against a reference fasta from Ensembl (CRCh38.101)

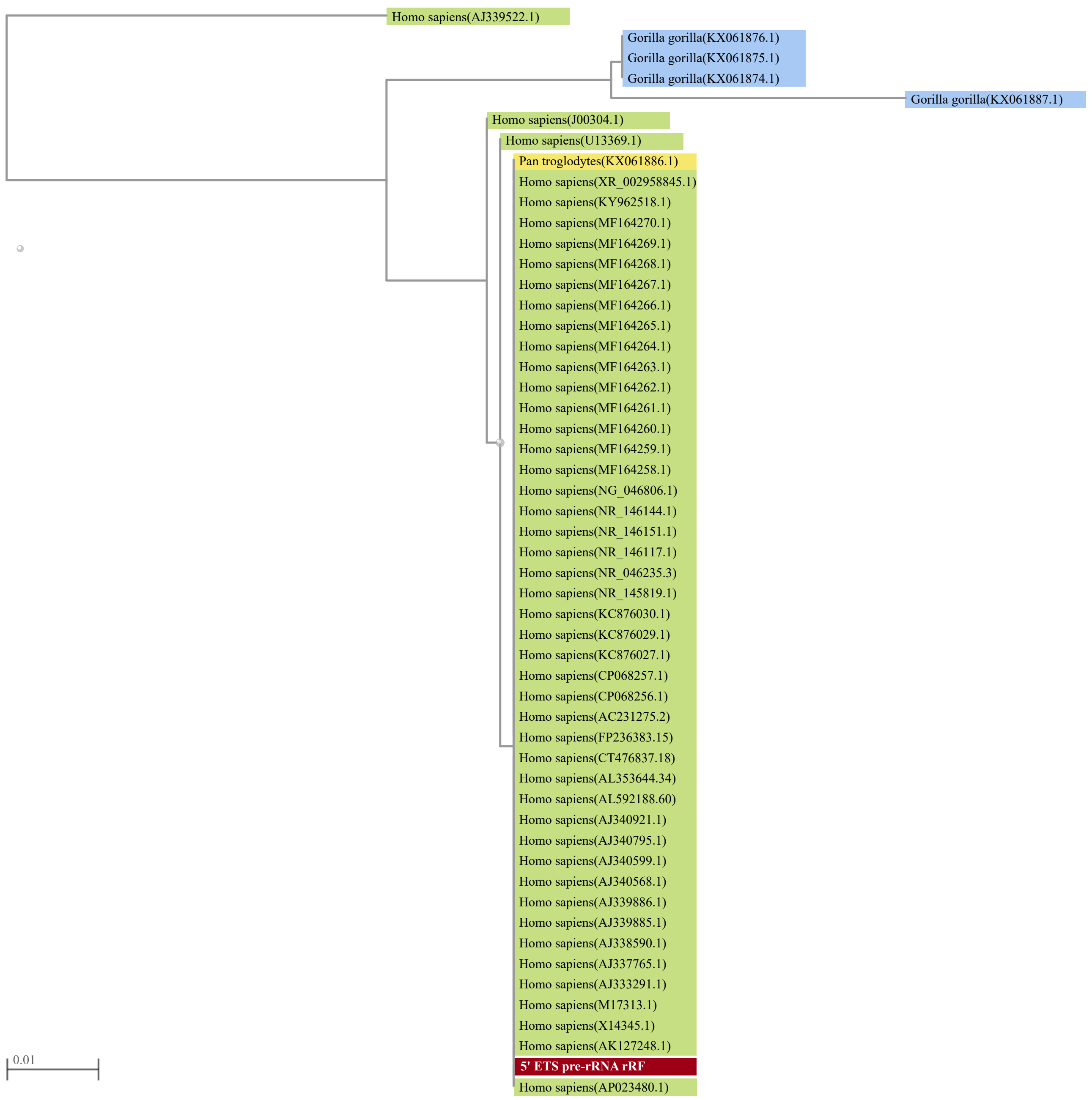


**Supplementary Figure S1. The 5' ETS pre-rRNA rRF was exclusive to african apes (Homininae).** Tree shows the distance in sequence similarity between the main fragment in Peak 1 (red) and the hits generated by aligning this sequence against multiple species at NCBI BLAST. Many highly similar matches were found in humans (green) and one sequence in chimpanzee (yellow), as well as a few similar sequences in gorillas (blue). The distance tree was constructed using fast minimum evolution (PMID: 14694080) with max sequence difference set to 0.75 (deafult at NCBI BLAST). The blasted (peak 1) seqeunce were: CTTCGTGATCGATGTGGTGACGTCGTGCTCTCCCGGGCCGGGTCCGAGCCGCGACGGGCGA

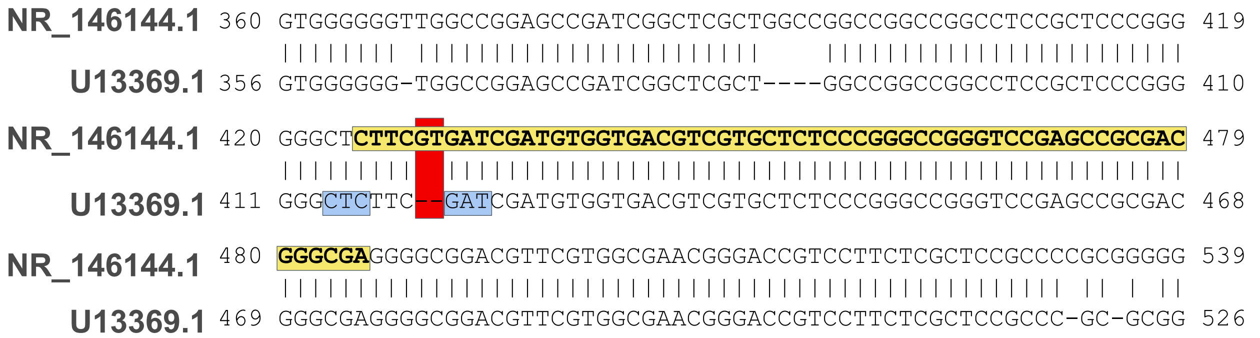


**Supplementary Figure S2. GT insertion is unique to 45S pre-rRNA that generates 5' ETS rRF generated from Peak 1.** Alignment between GenBank sequence NR_146144.1, which contains 5' ETS rRF from Peak 1, and U13369.1, which have been commonly used to map Human rRNA cleavage sites. Yellow and bold show the 5' ETS rRF from Peak 1. Blue shows where the C414-C416 and G420-U422 sites for 01 pre-rRNA cleavage in U13369.1 are located {Kass, 1987 #2548;Mullineux, 2012 #2547}. Red shows a G-T insertion in NR_146144.1, located right between C414-C416 and G420-U422.
