## Supplementary material for "Seqpac: A New Framework for small RNA analysis in R using Sequence-Based Counts": Supplementary_file_S1.R

```
#######################################################################################
#######################################################################################
#
# 2021-03-09, Linkoping University, Sweden. 
# Author: Daniel Natt
#
# These R scripts were published as a supplementary file to the original seqpac
# paper Skog et al. 2021. If using any of the scripts please site this paper.
#
# The latest version of seqpac can be downloaded from
# https://github.com/Danis102/seqpac Please, submit bug reports at github and
# specific questions at Biostar. Many questions can be answered by studying the
# package vignette and function manuals. For more important inquiries, please
# contact.
#
#
#######################################################################################
#######################################################################################


#######################################################################################
#####  1. Benchmarking seqpac make_trim function
#######################################################################################

# These scripts were used to generate some of the analysis reported in Fig 2 and 4
# We used a dataset originally published in:
# - Kang W, Eldfjell Y, Fromm B, Estivill X et al. miRTrace reveals the
# - organismal origins of microRNA sequencing data. Genome Biol 2018 Dec
# - 4;19(1):213. PMID: 30514392
# SRA project: SRP157338 
# ENA fastq download: https://www.ebi.ac.uk/ena/browser/view/PRJNA485638) 


#-------------------------------------------------------------------------------------
# Function to call cutadapt stand alone
#-------------------------------------------------------------------------------------
seqpac.make_cutadapt_alone <- function(input, output, parse=NULL, threads=1){
  
  ## Setup
  cat("\nRunning make_cutadapt ...")
  cat("\n--- cutadapt and fastq_quality_filter must be correctly installed.")
  if(!dir.exists(output)){ 
    suppressWarnings(dir.create(output)) 
  }
  fls <- list.files(input, pattern ="fastq.gz\\>|fastq\\>", full.names=TRUE, recursive=TRUE)
  
  # Make output names and check output folder
  nam <- gsub("\\.gz$", "", basename(fls))
  nam <- gsub("\\.fastq$|\\.fq$|fastq$", "", nam)
  nam <- gsub("\\.$|\\.tar$", "", nam)
  nam_trim <- paste0(nam, ".trim.fastq.gz")
  
  # Check for output folder
  out_file <- file.path(output, nam_trim)
  out_dir <- list.files(output, pattern=nam_trim, recursive = FALSE)
  if(length(out_dir)>0){
    stop(paste0("\n  Output trimmed fastq file names are identical to existing files in output:\n  ", out_dir, "\n  Please move or delete the file in the output folder:\n  ", output))
  }
  
  # cutadapt and fastq_quality_filter
  prog_report <- list(NULL) for(i in 1:length(fls)){
    log_lst <- list(NULL, NULL)
    names(log_lst) <- c("cutadapt", "fastq_quality_filter")
    spl_nam <- nam_trim[i]
    temp_out <- gsub("trim.fastq.gz$", "temp.fastq", out_file[i])
    if(!is.null(parse[[1]])){
      log_lst[[1]] <- system(paste0("cutadapt ", parse[[1]], " -o ", temp_out, " ", fls[i]), intern = TRUE)
    }
    if(!is.null(parse[[2]])){
      log_lst[[2]] <- system(paste0("fastq_quality_filter ", parse[[2]], " -v -i ", temp_out, " -o", out_file[i], " -z"), intern = TRUE)
    }
    if(!is.null(parse[[1]])){
      file.remove(temp_out) } prog_report[[1]] <- log_lst
  }
  doParallel::stopImplicitCluster()
  cat("\n--- Finished generating trimmed temporary files.")
  return(prog_report) gc(reset=TRUE) 
  }


#-------------------------------------------------------------------------------------
# Run benchmark
#-------------------------------------------------------------------------------------

# This function replicates adaptor trimming using make_trim, make_cutadapt and
# make_cutadapt_alone (above) 10 times. We ran it 6 times and saved time it took
# into a .csv text file with two columns named: "treatment" and "s" (=seconds).

# Note, Kang et al. uesd the TrueSeq small RNA kit, we therefore used the
# Illumina adaptor sequence for trimming.

library(rbenchmark) 
library(seqpac)

input = "<path to Kang et al. fastq files>"
output = "<temporary path for trimmed fastq files>"

bn <-
  rbenchmark::benchmark(
    #--------
    "seqpac" = {
      prog_report  <-  make_trim(input=input, output=output, threads=7, check_mem=FALSE, indels=TRUE, concat=12,
                                 adapt_3_set=c(type="hard_rm", min=10, mismatch=0.1),
                                 adapt_3="TGGAATTCTCGGGTGCCAAGGAACTCCAGTCACTTA",
                                 polyG=c(type="hard_trim",min=20, mismatch=0.1),
                                 seq_range=c(min=14, max=45), quality=c(threshold=20, percent=0.8))
      
      fn <- list.files(output, full.names=TRUE, recursive=FALSE)
      file.remove(fn) },
    #--------
    "cutadapt" = {
      prog_report  <-  make_cutadapt(input, output, threads=7,
                                     parse=list(cutadapt="-j 1 -a TGGAATTCTCGGGTGCCAAGGAACTCCAGTCACTTA --discard-untrimmed --nextseq-trim=20 -O 10 -m 14 -M 45",
                                                fastq_quality_filter="-q 20 -p 80"))
      fn <- list.files(output, full.names=TRUE, recursive=FALSE)
      file.remove(fn) },
    #--------
    "cutadapt_alone" = {
      prog_report  <- make_cutadapt_alone(input, output, threads=7,
                                          parse= list( cutadapt="-j 7 -a TGGAATTCTCGGGTGCCAAGGAACTCCAGTCACTTA --discard-untrimmed --nextseq-trim=20 -O 10 -m 14 -M 45",
                                                       fastq_quality_filter="-q 20 -p 80"))
      fn <- list.files(output, full.names=TRUE, recursive=FALSE)
      file.remove(fn) },
    #-------- 
    replications = 9, columns = c("test", "replications", "elapsed", "relative", "user.self", "sys.self", "user.child", "sys.child"))

# (replication = 9, the first cycle is not counted) 


#-------------------------------------------------------------------------------------
# Plot benchmark results (Fig 2A)
#-------------------------------------------------------------------------------------

library(ggplot2)
res_bench <- read.delim("<path to saved .csv file with benchmark results>", header=TRUE, sep=",")

ggplot(res_bench, aes(x=treatment, y=s, fill=treatment)) +
   							stat_summary(geom = "errorbar", fun.data = mean_se, position = "dodge") +
							stat_summary(geom = "bar", fun = mean, position = "dodge") +
							geom_jitter(col = "black", size=4, position=position_jitter(w=0.2, h=0.2)) +
                            scale_fill_manual(values=c("#094A6B", "grey", "blue"))+
							scale_color_manual(values=c("black", "black", "black"))+
							theme_bw()+
							theme(legend.position="none", axis.line.x =element_blank())


#######################################################################################
#####  2. Analyze PAC objects from Kang et al. 2018 dataset
#######################################################################################

# These scripts were used to generate some of the analysis reported in Fig 2 and 4
# We used a dataset originally published in:
#
#   Kang W, Eldfjell Y, Fromm B, Estivill X et al. miRTrace reveals the
#   organismal origins of microRNA sequencing data. Genome Biol 2018 Dec
#   4;19(1):213. PMID: 30514392
#
# SRA project: SRP157338 
# ENA fastq download: https://www.ebi.ac.uk/ena/browser/view/PRJNA485638) 


#-------------------------------------------------------------------------------------
# Generate PAC objects for Kang et al. (Fig 2B/C)
#-------------------------------------------------------------------------------------

# For generating PAC objects for some of the analysis in Fig 2 we used the
# make_trim (internal) and make_cutadapt (external) functions in seqpac.

input = "<path to Kang et al. fastq files>"
pheno_input = "<path to unpacked: Kang_pheno.csv>" # In Supplementary_file_S3.zip

# Note: pheno.csv was manually constructed from SRA metadata text files. See
# vignette for more details on how to make a pheno input file.

#--------
# Using make_trim (seqpac):
parse = list(adapt_3_set=c(type="hard_rm", min=10, mismatch=0.1),
             adapt_3="TGGAATTCTCGGGTGCCAAGGAACTCCAGTCAC",
             polyG=c(type="hard_trim", min=20, mismatch=0.1),
             seq_range=c(min=14, max=45), quality=c(threshold=20, percent=0.8),
             indels=TRUE, concat=12, check_mem=FALSE)
counts_seq  <- make_counts(input, threads=7, parse=parse, type="fastq",
                           trimming="seqpac", plot=TRUE, evidence=c(experiment=2, sample=1))
anno_seq <- make_anno(counts_seq, type="counts")
pheno_seq <- make_pheno(pheno_input = pheno_input,
                        type="manual", counts=counts_seq[[1]], progress_report=counts_seq[[2]])

pac_seq <- make_PAC(pheno_input=pheno_input, anno_input=anno_seq, counts_input=counts_seq[[1]]) 
PAC_check(pac_seq)
# save(pac_seq, file="<output path/pac_master_seqpac.Rdata>")

#--------
# Using make_cutadapt (cutadapt):
parse = list(cutadapt="-j 1 -a TGGAATTCTCGGGTGCCAAGGAACTCCAGTCAC --discard-untrimmed --nextseq-trim=20 -O 10 -m 14 -M 45 -e 0.1",
             fastq_quality_filter="-q 20 -p 80")
counts_cut  <- make_counts(input, threads=7, parse=parse, type="fastq", trimming="cutadapt", plot=TRUE, evidence=c(experiment=2, sample=1))
anno_cut  <- make_anno(counts_cut, type="counts")
pheno_cut <- make_pheno(pheno_input=pheno_input,
                        type="manual", counts=counts_cut[[1]], progress_report=counts_cut[[2]])

pac_cut <- make_PAC(pheno_input=pheno_cut, anno_input=anno_cut, counts_input=counts_cut[[1]]) 
PAC_check(pac_cut)
# save(pac_cut, file="<output path/pac_master_cutadapt.Rdata>")


#-------------------------------------------------------------------------------------
# Reannotation workflow for Kang et al. 2018   
#-------------------------------------------------------------------------------------

# Only the PAC object generated internally in seqpac (using the make_trim function)
# was annotated using the reannotation workflow.

load("<output path/pac_master_seqpac.Rdata>")

output_path = "<path to output folder>"

#--------
# Genome alignment:

ref_paths <- list(dm6="<path to bowtie indexed dm6 fruit fly fasta genome filer>",
                      hg38="<path to bowtie indexed hg38 human fasta genome file>")

map_reanno(pac_seq, type = "internal", output_path, ref_paths = ref_paths,
           mismatches = 3, threads = 7, import="genome")
reanno_seq <- make_reanno(reanno_path=output_path, PAC=pac_seq, mis_fasta_check = TRUE, threads = 7)
pac_seq <- add_reanno(reanno=reanno_seq, mismatches = 3,  merge_pac=pac_seq, type = "genome", genome_max = 10)
# save(pac_seq, file="<output path/pac_master_seqpac.Rdata>")

# Sequences in total: 491968 #
# mis0	dm6:  	Ref_hits= 57901     (11.77%)
# mis0	hg38: 	Ref_hits= 229082    (46.56%)
# mis1	dm6:  	Ref_hits= 22014     (4.47%)
# mis1	hg38: 	Ref_hits= 88590     (18.01%)
# mis2	dm6:  	Ref_hits= 13616     (2.77%)
# mis2	hg38: 	Ref_hits= 54194     (11.02%)
# mis3	dm6:    Ref_hits= 15112     (3.07%)
# mis3	hg38: 	Ref_hits= 36839     (7.49%)


#--------
# sRNA class annotations

ref_paths_bio <- list(dm6_miRNA="<path to fruit fly miRBase fasta: miRBase_21-dme.fa>",
                      dm6_Ensembl="<path to fruit fly Ensembl ncRNA fasta: Drosophila_melanogaster.BDGP6.ncrna.fa>",
                      dm6_tRNA="<path to fruit fly GtRNAdb fasta: dm6-tRNAs_mature.fa>",
                      dm6_piRNA="<path to fruit fly pirBase fasta: piR_dme.fa",
                      hg38_miRNA="<path to human miRBase fasta: miRBase_21-hsa.fa>",
                      hg38_Ensembl="<path to human Ensembl ncRNA fasta: Homo_sapiens.GRCh38.ncrna.fa>",
                      hg38_tRNA="<path to human GtRNAdb fasta: hg38-tRNAs_mature.fa>",
                      hg38_piRNA="<path to human pirBase fasta: piR_hsa.fa")

bio_search <- list( dm6_Ensembl=c("lincRNA", "miRNA", "rRNA", "pre_miRNA", "snoRNA", "snRNA", "_tRNA", "mt_tRNA"),
                    dm6_miRNA="dme-",
                    dm6_piRNA = "piR-dme|piRNA",
                    dm6_tRNA =c("^tRNA", "mt-tRNA"),
                    hg38_miRNA="hsa",
                    hg38_Ensembl=c("lncRNA", "miRNA", "rRNA", "snoRNA", "snRNA", "tRNA"),
                    hg38_tRNA=c("tRNA"),
                    hg38_piRNA="piR-hsa|piRNA")

hierarchy <- list(rRNA="rRNA", tRNA="tRNA", miRNA ="miRNA", snoRNA="snoRNA", snRNA="snRNA", lncRNA="lncRNA|lincRNA", piRNA="piRNA" )

map_reanno(pac_seq, type = "internal", output_path, ref_paths=ref_paths_bio, mismatches = 3, threads = 7, import="biotype")
reanno_seq <- make_reanno(reanno_path=output_path, PAC=pac_seq, mis_fasta_check = TRUE, threads = 7)
pac_seq <- add_reanno(reanno_seq, bio_search=bio_search, type="biotype", bio_perfect=FALSE, mismatches = 0, merge_pac=pac_seq)
pac_seq <- simplify_reanno(pac_seq, hierarchy=hierarchy, mismatches = 0, bio_name = "Biotype", merge_pac = TRUE)
#save(pac_seq, file="<output path/pac_master_seqpac.Rdata>")


#-------------------------------------------------------------------------------------
# Reannotation plots using the Kang et al. 2018 dataset (Fig 4A/B/C)  
#-------------------------------------------------------------------------------------
load("<output path/pac_master_seqpac.Rdata>")

#--------
# Species genome mapping pie (Fig 4A)
pac_seq$Anno$mis0_genome_olap <- paste(ifelse(pac_seq$Anno$dm6 == "mis0", "fly", "no"), ifelse(pac_seq$Anno$hg38 == "mis0", "human", "no"))
plts_mis0 <- PAC_pie(pac_seq, pheno_target=list("Sample_ID"), anno_target=list("mis0_genome_olap"))
cowplot::plot_grid(plotlist=plts_mis0, nrow=3, ncol=3, scale=0.8)

pac_seq$Anno$mis3_genome_olap <- paste(ifelse(pac_seq$Anno$dm6 %in% c("mis0","mis1", "mis2", "mis3"), "fly", "no"),
                                       ifelse(pac_seq$Anno$hg38 %in% c("mis0","mis1", "mis2", "mis3"), "human", "no"))

plts_mis3 <- PAC_pie(pac_seq, pheno_target=list("Sample_ID"),
                     anno_target=list("mis3_genome_olap", c("no no", "fly no", "no human", "fly human")))

cowplot::plot_grid(plotlist=plts_mis3, nrow=3, ncol=3, scale=0.8)


#--------
# Size dist with classes (Fig 4B)
ord <- c("no_anno", "other", "miRNA", "tRNA", "snoRNA", "piRNA" , "snRNA", "lncRNA", "rRNA")
sizebio_seq <- PAC_sizedist(pac_seq, anno_target=list("Biotype", ord))
cowplot::plot_grid(plotlist=sizebio_seq$Histograms)

#--------
# Nbias plots (Fig 4C)
nbias_seq <- PAC_nbias(pac_seq)
cowplot::plot_grid(plotlist=nbias_seq$Histograms)


#######################################################################################
#####  2. Detecting contamination in Tong et al. 2020 dataset (cancer cell lines)
#######################################################################################

# These scripts were used to generate some of the analysis reported in Fig 7.
# We used a dataset originally published in:
#
#   Tong F, Andress A, Tang G, Liu P et al. Comprehensive profiling of
#   extracellular RNA in HPV-induced cancers using an improved pipeline for small
#   RNA-seq analysis. Sci Rep 2020 Nov 10;10(1):19450. PMID: 33173155
# 
# SRA project: SRP285629 
# ENA fastq download: https://www.ebi.ac.uk/ena/browser/view/PRJNA666144) 


#-------------------------------------------------------------------------------------
# PAC generation Tong et al. 2020 dataset
#-------------------------------------------------------------------------------------

# Since Tong et al. 2020 used an Illumina HiSeq 3000 sequencer with a flow cell
# kit resulting in 50 nt reads, we ran 'make_trim' with 'type="hard_trim"' in
# 'adapt_3_set' input. This will trim 3' adaptor sequence, and if no adaptor
# sequence is found, the read will still be saved. In the Kang et al. 2018 and
# Skog et al. 2021 datasets, which used Illumina NextSeq 500 sequencers with
# kits generating 75 nt long reads, we set 'type="hard_rm"'. With longer reads
# this option is the best, since it will remove reads that fail to contain
# adaptor sequence, which guarantees that all sequences came from short sRNA.
# For the Tong et al 2020 dataset we also set the maximum read lenght > 50 nt to
# investigate the ratios of long RNA in this dataset.

# Note, the 'cell_type' and 'type' columns in the pheno.csv was obtained from
# the 'sample_title' column of the ENA report available at
# https://www.ebi.ac.uk/ena/browser/view/PRJNA666144)


path_to_fastq <- "<your path to downloaded fastq files>"
pheno_path  = "<path to unpacked: Tong_pheno.csv>" # In Supplementary_file_S3.zip

parse <- list(polyG=c(type="hard_trim", min=20, mismatch=0.1),
                        adapt_3_set=c(type="hard_trim", min=10, mismatch=0.1),
                        adapt_3="AGATCGGAAGAGCACACGTCTGAACTCCA",
                        seq_range=c(min=14, max=70),
                        quality=c(threshold=20, percent=0.8),
                        indels=TRUE, concat=12, check_mem=FALSE)
count_list <- make_counts(input=path_to_fastq, type = "fastq", trimming="seqpac", 
                          parse=parse, threads=12, save_temp = TRUE)
anno <- make_anno(count_list, type = "counts", stat = TRUE)

pheno <- make_pheno(pheno_input=pheno_path,
                    counts=count_list$counts,
                    progress_report=count_list$progress_report)
pac  <- make_PAC(pheno_input=pheno, anno_input=anno, counts_input=count_list$counts)


#-------------------------------------------------------------------------------------
# Principle component analysis on filtered data (Fig 7A) 
#-------------------------------------------------------------------------------------
# Note, enteries in cell_type column:
pheno_ord <- c("C33A", "C41", "Caski", "HeLa", "HT-3", "SiHa", "SW756", "SCC-090", "SCC-1", "SCC-47", "UPCI-017", "UPCI-068", "SCC-154", "SCC-4")

pac_filt  <- PAC_filter(pac, size = c(16,50), threshold=10, coverage=50, pheno_target=list("cell_type", pheno_ord), stat=TRUE)
pac_filt  <- PAC_norm(pac_filt, norm = "vst")
pac_filt  <- PAC_norm(pac_filt, norm = "cpm")

pca_out <- PAC_pca(pac_filt, norm="vst", pheno_target=list("type"))


#-------------------------------------------------------------------------------------
# Reannotation workflow for filtered Tong et al. 2020 dataset
#-------------------------------------------------------------------------------------

# Notice, for running the reannotation workflow you need bowtie index fasta
# reference files. Here is an example on how this can be done using the Rbowtie
# package:

# Rbowtie::bowtie_build("<your path to bowtie indexed Mycoplasma hyorhinis fasta/GCF_000383515.1_ASM38351v1_genomic.fa", 
#                       outdir="<same folder as fasta>",
#                       prefix="<fasta file name without file extension, like: GCF_000383515.1_ASM38351v1_genomic", 
#                       force=TRUE)

#--------
# Genome alignment
output_path = "<your path to working directory for bowtie>"
ref_paths <- list(hg38="<your path to bowtie indexed human genome/Homo_sapiens.GRCh38.dna.toplevel.fa",
                      M.hyorhinis="<your path to bowtie indexed Mycoplasma hyorhinis fasta/GCF_000383515.1_ASM38351v1_genomic.fa")

map_reanno(pac_filt, type = "internal", output_path, ref_paths = ref_paths,
           mismatches = 3, threads = 12, import="genome")
reanno_seq <- make_reanno(reanno_path=output_path, PAC=pac_filt, mis_fasta_check = TRUE)
pac_filt <- add_reanno(reanno=reanno_seq, mismatches = 3,  merge_pac=pac_filt, type = "genome", genome_max = 10)


#--------
# sRNA class
ref_paths_bio <- list(hg38_miRNA="<path to human miRBase fasta: miRBase_21-hsa.fa>",
                      hg38_Ensembl="<path to human Ensembl ncRNA fasta: Homo_sapiens.GRCh38.ncrna.fa>",
                      hg38_tRNA="<path to human GtRNAdb fasta: hg38-tRNAs_mature.fa>",
                      hg38_piRNA="<path to human pirBase fasta: piR_hsa.fa")

bio_search <- list( hg38_miRNA="hsa",
                    hg38_Ensembl=c("lncRNA", "miRNA", "rRNA", "snoRNA", "snRNA", "tRNA"),
                    hg38_tRNA=c("tRNA"),
                    hg38_piRNA="piR-hsa|piRNA")

hierarchy <- list(rRNA="rRNA", tRNA="tRNA", miRNA ="miRNA", snoRNA="snoRNA", snRNA="snRNA", lncRNA="lncRNA|lincRNA", piRNA="piRNA" )

map_reanno(pac_filt, type = "internal", output_path, ref_paths=ref_paths_bio, mismatches = 3, threads = 12, import="biotype")
reanno_seq <- make_reanno(reanno_path=output_path, PAC=pac_filt, mis_fasta_check = TRUE)

pac_filt <- add_reanno(reanno_seq, bio_search=bio_search, type="biotype", bio_perfect=FALSE, mismatches = 3, merge_pac=pac_filt)
pac_filt <- simplify_reanno(pac_filt, hierarchy=hierarchy, mismatches = 0, bio_name = "Biotype_mis0", merge_pac = TRUE)
pac_filt <- simplify_reanno(pac_filt, hierarchy=hierarchy, mismatches = 3, bio_name = "Biotype_mis3", merge_pac = TRUE)


#-------------------------------------------------------------------------------------
# Better rRNA classification using PAC_gtf
#-------------------------------------------------------------------------------------
# NCBI refseq has sometimes better coverage of rRNA like sequences than Ensembl
# ncRNA. The latest refseq coordinates for the human genome can be found here:
# https://ftp.ncbi.nlm.nih.gov/refseq/H_sapiens/annotation/GRCh38_latest/refseq_identifiers/

#--------
# Prepare rRNA exclusive GTF:
gtf <- tibble::as_tibble(rtracklayer::readGFF("<your path to gtf/GRCh38_latest_genomic.gtf>"))
gtf_genes <- gtf[gtf$type == "gene",]
gtf_rrna <- gtf_genes[grepl("^RNA", gtf_genes$description),]
gtf_rrna <- gtf_rrna[grepl("ribosomal", gtf_rrna$description),]
table(gtf$gene_biotype)
table(gtf_genes$gene_biotype)
table(gtf_rrna$gene_biotype)
rtracklayer::export(gtf_rrna, "<your path to gtf/rRNA_extract.gtf", format="gtf")

# Important, you may need to convert the chromosomal names of NCBI/refseq so
# that they match with the genome alignments generated in the reannotation
# workflow (Ensembl). Devon Ryan's github page contains nice tables for converting
# chromosomal named between databases:
# https://github.com/dpryan79/ChromosomeMappings

#--------
# Run PAC_gtf allowing for 0 and 3 mismatches:
gtf_rrna <- tibble::as_tibble(rtracklayer::readGFF("<your path to gtf/rRNA_extract.gtf"))
gtf_anno_mis0 <- PAC_gtf(PAC=pac_filt, genome="<your path to bowtie indexed human genome/Homo_sapiens.GRCh38.dna.toplevel.fa", 
                         gtf_other=list(rrna=gtf_rrna), 
                         target_other=list(rrna=c("gene", "gene_id", "gene_biotype", "description")), 
                         mismatches=0, threads=12, return="simplify")

gtf_anno_mis3 <- PAC_gtf(PAC=pac_filt, genome="/data/Data_analysis/Genomes/Humans/Ensembl/CRCh38.101/Homo_sapiens.GRCh38.dna.toplevel.fa", 
                         gtf_other=list(rrna=gtf_rrna), 
                         target_other=list(rrna=c("gene", "gene_id", "gene_biotype", "description")), 
                         mismatches=3, threads=12, return="simplify" )

#--------
# Update new biotype mis variable and then simplify with hierarchry
new_var_mis0 <- paste(pac_filt$Anno$mis0_bio, ifelse(!is.na(gtf_anno_mis0$gene_biotype), "hg38_refSeq_rRNA", ""), sep=";")
pac_filt$Anno$mis0_bio <- gsub(";$", "", new_var_mis0)
hierarchy <- list(rRNA="rRNA", tRNA="tRNA", miRNA ="miRNA", snoRNA="snoRNA", snRNA="snRNA", lncRNA="lncRNA|lincRNA", piRNA="piRNA" )
pac_filt <- simplify_reanno(pac_filt, hierarchy=hierarchy, mismatches = 0, bio_name = "Biotype_mis0_refseq", merge_pac = TRUE)


#-------------------------------------------------------------------------------------
# Make sRNA class sizedist (Fig 7B) 
#-------------------------------------------------------------------------------------

#--------
# Fix subgroups
pac_filt$Pheno$subtype <- ifelse(pac_filt$Pheno$cell_type %in% "SCC-154", "SCC-154",
                                 ifelse(pac_filt$Pheno$cell_type %in% "SCC-4", "SCC-4", "all_other"))
pac_filt$Pheno$subtype2 <- paste(pac_filt$Pheno$subtype, pac_filt$Pheno$type, sep="_")

#--------
# Mixing means of all with individual samples of SCC4 and SCC154
pac_filt <- PAC_summary(pac_filt, norm = "cpm", type = "means", pheno_target=list("subtype2", c("all_other_cell", "all_other_EXO", "all_other_MV")), merge_pac = TRUE)
temp <-  pac_filt$norm$cpm[,rownames(pac_filt$Pheno)[!grepl("all", pac_filt$Pheno$subtype2)]]
colnames(temp) <- pac_filt$Pheno$subtype2[!grepl("all", pac_filt$Pheno$subtype2)]
pac_filt$summary$cpmMeans_subtype2 <- cbind(pac_filt$summary$cpmMeans_subtype2, temp)

#--------
# plot size dist for subtype 2
ord <-  c("no_anno", "other",  "snoRNA",  "snRNA", "rRNA", "tRNA", "piRNA",  "miRNA")
out_sizedist_mis0 <- PAC_sizedist(pac_filt, norm = NULL, anno_target= list("Biotype_mis0_refseq", ord), summary_target=list("cpmMeans_subtype2"))
cowplot::plot_grid(plotlist=out_sizedist_mis0$Histograms)


#-------------------------------------------------------------------------------------
# Plot percent mycoplasma (Fig 7C) 
#-------------------------------------------------------------------------------------

pac_filt$Anno$genome_hit <- ifelse(grepl("mis",  pac_filt$Anno$hg38_genome), "human",
                              ifelse(grepl("mis",  pac_filt$Anno$M.hyorhinis_genome), "mycoplasma", "no_hit"))
ord <-  c("no_hit", "mycoplasma", "human")
PAC_stackbar(pac_filt, anno_target= list("genome_hit", ord),  pheno_target=list("type"), color=c("#BCBCBD", "#9D0014","#BCD771"), total = FALSE)


#-------------------------------------------------------------------------------------
# Principle component analysis after non-human removal (Fig 7D) 
#-------------------------------------------------------------------------------------

#--------
# Make a pure human filter
pure_filt <- 
  as.data.frame(
    cbind(grepl("mis",  pac_filt$Anno$hg38_genome),
          grepl("mis",  pac_filt$Anno$M.hyorhinis_genome)))

pure_filt <- apply(pure_filt, 1,  function(x){paste(x, collapse="_")}) == "TRUE_FALSE"
pac_filt$Anno$pure<- ifelse(pure_filt==TRUE, "pure_human", "not")


#--------
# IMPORTANT: Save PAC object for pre-rRNA analysis (next section)
save(pac_filt, file="<your path to save folder/pac_filt_Tong2020.Rdata>")

#--------
# Filter, remove long fragments, and add new vst normalization
pac_filt  <- PAC_filter(pac_filt, size = c(16,45), anno_target=list("pure", "pure_human"))
pac_filt$norm <- pac_filt$norm[!names(pac_filt$norm) %in% "vst"]
pac_filt <- PAC_norm(pac_filt, norm = "vst")

#--------
# Run PCA (Fig 7D)
pca_out <- PAC_pca(pac_filt, norm="vst", pheno_target=list("type"), graphs=TRUE)


#######################################################################################
#####  3. Novel 5' ETS pre-rRNA rRF in Skog et al. 2021 (BMH21 exposure in HeLA)  
#######################################################################################       

# These scripts were used to generate some of the analysis reported in Fig 8.
# We primarily used a dataset generated within the present study (Skog et al. 2021):
# SRA project: SRP309937 
# ENA fastq download: https://www.ebi.ac.uk/ena/browser/view/PRJNA708219) 

#-------------------------------------------------------------------------------------
# PAC generation Skog et al. 2021 dataset 
#-------------------------------------------------------------------------------------

# Important, in the SRA metadata the 'sample_name' column holds the different
# treatment groups. Since we had 75 nt reads, we used the "hard_rm" option in
# the 'adapt_3_set' setting, removing reads that did not contain the NEBNext
# adaptor.

input <- "<your path to downloaded fastq files>"
pheno_path  = "<path to unpacked: Skog_pheno.csv>" # In Supplementary_file_S3.zip

parse <- list(polyG=c(type="hard_trim", min=20, mismatch=0.1),
                     adapt_3_set=c(type="hard_rm", min=10, mismatch=0.1), 
                     adapt_3="AGATCGGAAGAGCACACGTCTGAACTCCA",
                     seq_range=c(min=14, max=70),
                     quality=c(threshold=20, percent=0.8),
                     indels=TRUE, concat=12, check_mem=FALSE)

count_list <- make_counts(input=input, type = "fastq", trimming="seqpac", 
                          parse=parse, threads=12, save_temp = TRUE)
anno <- make_anno(count_list, type = "counts", stat = TRUE)
pheno <- make_pheno(pheno_input=pheno_path,
                    counts=count_list$counts,
                    progress_report=count_list$progress_report)
#--------
# Fix shorter group names
pheno$treatment <- ifelse(grepl("DMSO", pheno$sample_name),  "DMSO_60",
                          ifelse(grepl("60 min BMH21", pheno$sample_name), "BMH21_60", 
                            ifelse(grepl("12h BMH21", pheno$sample_name), "BMH21_12", "bad")))


pac  <- make_PAC(pheno_input=pheno, anno_input=anno, counts_input=count_list$counts)
table(pac$Pheno$treatment)


#-------------------------------------------------------------------------------------
# PAC filtering, reannotation and normalization
#-------------------------------------------------------------------------------------

# Here, we used a slightly more advanced normalization and filtering method.
# This is done in five steps: 
# 1. Low-level filtering to get rid of noise prior to reannoation
# 2. Multi-genome alignment (reannotation workflow)
# 3. Removal of non-human/mycoplasmic sequences
# 4. Normalization (cpm and vst) 
# 5. High-level filtering to obtain the most relevant sRNA

# We also used the PAC_filtsep function to filter within treatment groups, as
# described in the vignette and main text of the article.

#--------
# Low-level filtering (at least 3 counts within a treatment group, ranging between 16-65 nt) 
filtsep <- PAC_filtsep(pac, norm = "counts", threshold = 3, coverage = 100,
                       pheno_target=list("treatment"))
pac_filt_low  <- PAC_filter(pac, size = c(16,65), anno_target=unique(unlist(as.list(filtsep))))


#--------
# Multi-genome alignment
output_path = "<your path to working directory for bowtie>"
ref_paths <- list(hg38="<your path to bowtie indexed human genome/Homo_sapiens.GRCh38.dna.toplevel.fa",
                  M.hyorhinis="<your path to bowtie indexed Mycoplasma hyorhinis fasta/GCF_000383515.1_ASM38351v1_genomic.fa")

map_reanno(pac_filt_low, type = "internal", output_path, ref_paths = ref_paths,
           mismatches = 3, threads = 12, import="genome")
reanno_seq <- make_reanno(reanno_path=output_path, PAC=pac_filt_low, mis_fasta_check = TRUE)
pac_filt_low <- add_reanno(reanno=reanno_seq, mismatches = 3,  merge_pac=pac_filt_low, type = "genome", genome_max = 10)


#--------
# Removal of mycoplasmic sequences
pac_filt_low  <- PAC_filter(pac_filt_low, size = c(16,65), anno_target=list("M.hyorhinis_genome", "_"))

#--------
# Normalization
pac_filt_low  <- PAC_norm(pac_filt_low, norm = "cpm")
pac_filt_low  <- PAC_norm(pac_filt_low, norm = "vst")

#--------
# High-level filtering
filtsep <- PAC_filtsep(pac_filt_low, norm = "cpm", threshold = 10, coverage = 100,
                       pheno_target=list("treatment"))
pac_filt  <- PAC_filter(pac_filt_low, size = c(16,65), anno_target=unique(unlist(as.list(filtsep))))


#-------------------------------------------------------------------------------------
# Reanno workflow sRNA class
#-------------------------------------------------------------------------------------
# We already did the genome alignment in the previous step, know we do the sRNA
# class alignments

#--------
# sRNA class
ref_paths_bio <- list(hg38_miRNA="<path to human miRBase fasta: miRBase_21-hsa.fa>",
                      hg38_Ensembl="<path to human Ensembl ncRNA fasta: Homo_sapiens.GRCh38.ncrna.fa>",
                      hg38_tRNA="<path to human GtRNAdb fasta: hg38-tRNAs_mature.fa>",
                      hg38_piRNA="<path to human pirBase fasta: piR_hsa.fa")

bio_search <- list( hg38_miRNA="hsa",
                    hg38_Ensembl=c("lncRNA", "miRNA", "rRNA", "snoRNA", "snRNA", "tRNA"),
                    hg38_tRNA=c("tRNA"),
                    hg38_piRNA="piR-hsa|piRNA")

hierarchy <- list(rRNA="rRNA", tRNA="tRNA", miRNA ="miRNA", snoRNA="snoRNA", snRNA="snRNA", lncRNA="lncRNA|lincRNA", piRNA="piRNA" )

map_reanno(pac_filt, type = "internal", output_path, ref_paths=ref_paths_bio, mismatches = 3, threads = 12, import="biotype")
reanno_seq <- make_reanno(reanno_path=output_path, PAC=pac_filt, mis_fasta_check = TRUE)

pac_filt <- add_reanno(reanno_seq, bio_search=bio_search, type="biotype", bio_perfect=FALSE, mismatches = 3, merge_pac=pac_filt)
pac_filt <- simplify_reanno(pac_filt, hierarchy=hierarchy, mismatches = 0, bio_name = "Biotype_mis0", merge_pac = TRUE)
pac_filt <- simplify_reanno(pac_filt, hierarchy=hierarchy, mismatches = 3, bio_name = "Biotype_mis3", merge_pac = TRUE)


#--------
# Add rRNA classification using PAC_gtf (see above section 2)
gtf_rrna <- tibble::as_tibble(rtracklayer::readGFF("<your path to gtf/rRNA_extract.gtf"))
gtf_anno_mis0 <- PAC_gtf(PAC=pac_filt, genome="<your path to bowtie indexed human genome/Homo_sapiens.GRCh38.dna.toplevel.fa>", 
                         gtf_other=list(rrna=gtf_rrna), 
                         target_other=list(rrna=c("gene", "gene_id", "gene_biotype", "description")), 
                         mismatches=0, threads=12, return="simplify")

gtf_anno_mis3 <- PAC_gtf(PAC=pac_filt, genome="<your path to bowtie indexed human genome/Homo_sapiens.GRCh38.dna.toplevel.fa>", 
                         gtf_other=list(rrna=gtf_rrna), 
                         target_other=list(rrna=c("gene", "gene_id", "gene_biotype", "description")), 
                         mismatches=3, threads=12, return="simplify" )

new_var_mis0 <- paste(pac_filt$Anno$mis0_bio, ifelse(!is.na(gtf_anno_mis0$gene_biotype), "hg38_refSeq_rRNA", ""), sep=";")
pac_filt$Anno$mis0_bio <- gsub(";$", "", new_var_mis0)
hierarchy <- list(rRNA="rRNA", tRNA="tRNA", miRNA ="miRNA", snoRNA="snoRNA", snRNA="snRNA", lncRNA="lncRNA|lincRNA", piRNA="piRNA" )
pac_filt <- simplify_reanno(pac_filt, hierarchy=hierarchy, mismatches = 0, bio_name = "Biotype_mis0_refseq", merge_pac = TRUE)


#-------------------------------------------------------------------------------------
# Skog et al 2021 DE analysis using PAC_deseq (Fig 8A; Supplementary Table 1)
#-------------------------------------------------------------------------------------
# Our hypothesis was that sRNA classified as being derived from rRNA (rRFs)
# should be affected by RNA pol I inhibition for 60 min (anticancer substance:
# BMH21). Thus we extracted sRNA originating from rRNA and performed an
# differental expression ananlysis. Since many rRFs are highly expressed, we
# only targeted highly expressed rRFs (>=100 cpm within a group) to avoid
# problems with false negatives following multiple testing.

#--------
## Extract the BMH21 60 min and DMSO 60 min groups and filter on high expression
pac_short <- PAC_filter(pac_filt_hi, pheno_target=list("treatment", c("DMSO_60", "BMH21_60")))
filtsep <- PAC_filtsep(pac_short, norm = "cpm", threshold = 100, coverage = 100, pheno_target=list("treatment"))
pac_short <- PAC_filter(pac_short, anno_target=unique(unlist(filtsep))) 

#--------
# Extract likely rRFs and perform DE analysis 
pac_short_rRNA <- PAC_filter(pac_short, anno_target=list("Biotype_mis3_refseq", "rRNA"))
deseq_out_rRNA <- PAC_deseq(pac_short_rRNA, model=~treatment, pheno_target=list("treatment", c("BMH21_60", "DMSO_60")))
ord_res_rRNA <- deseq_out_rRNA$result[order(deseq_out_rRNA$result$padj),]
ord_res_rRNA[1:21, c(16, 2:3,7, 18)]
# Highest expression (baseMean:  6240.5183; Size 61 nt:
# CTTCGTGATCGATGTGGTGACGTCGTGCTCTCCCGGGCCGGGTCCGAGCCGCGACGGGCGA (in Fig 8K) 


#-------------------------------------------------------------------------------------
# Coverage plot of pre-rRNA Skog et al. 2021  (present study: Fig 8B-F) 
#-------------------------------------------------------------------------------------
# In 'Supplementary_file_2.fa' that are published with the article are the fasta
# references sequences used for generating Fig 8B-L. It includes the following
# reference sequences:
#
# "NR_146144.1"                                (downloaded from NCBI nucleotide)                               
# "Peak1_hg38_dna_range=chr21:8206319-8206669" (downloaded from UCSC genome browser)
# "Peak2_hg38_dna_range=chr21:8212475-8212825" (downloaded from UCSC genome browser)
# "Peak3_hg38_dna_range=chr21:8213765-8214115" (downloaded from UCSC genome browser)
# "Peak4_hg38_dna_range=chr21:8218787-8219137" (downloaded from UCSC genome browser)
# "47S_U13369.1"                               (downloaded from NCBI nucleotide)     
#
# Graphs are generated by running the PAC_mapper in combination with the
# PAC_covplot functions, as follows:

#--------
# Make treatment means and run PAC_mapper
pac_filt_hi <- PAC_summary(pac_filt_hi, norm="cpm", pheno_target=list("treatment"))

ref <- "<your path to Supplementary_file_2.fa>"
map_object <- PAC_mapper(pac_filt_hi, ref=ref, N_up = "NNN", N_down = "NNN", mapper="reanno", mismatches=0, threads=8, report_string=FALSE)
out_covp <- PAC_covplot(pac_filt_hi, map=map_object, summary_target= list("cpmMeans_treatment", c("DMSO_60", "BMH21_60")), xseq = FALSE)

out_covp[1] # High peak in 5' ETS region
cowplot::plot_grid(plotlist=out_covp[2:5], nrow=4, ncol=1) # Lower peak in 5' ETS region
out_covp[6] # Lower peak in 5' ETS region


#-------------------------------------------------------------------------------------
# Plots error bars for rRF peaks Skog et al. 2021  (Fig 8G-J) 
#-------------------------------------------------------------------------------------
# The map object generated by PAC_mapper can be used to extract fragments
# aligning to the same transcript, as follows:

#--------
# Create new Peak variable in PAC object

# First extract peak sequences from PAC object: 
logi_peak1<- rownames(pac_filt_hi$Anno) %in% rownames(map_object[[2]]$Alignments)
logi_peak2<- rownames(pac_filt_hi$Anno) %in% rownames(map_object[[3]]$Alignments)
logi_peak4 <- rownames(pac_filt_hi$Anno) %in% rownames(map_object[[5]]$Alignments)

# Note, Peak 3 seem to have two main fragments, one shorter (Peak 3a) and one longer. From
# the coverage plot (Fig 8E) we can see that the longest Peak 3a fragment is
# "CGCGACCTCAGATCAGACGTG". We use a substring, "GCGACCTCAGA", of this main Peak
# 3a fragment to 'catch' additional Peak 3a fragments, then we extracted only
# fragments shorter than the longest peak (<=24 nt).

logi_peak3_1 <- rownames(pac_filt_hi$Anno) %in% rownames(map_object[[4]]$Alignments)
logi_peak3_2 <- grepl("GCGACCTCAGA", rownames(pac_filt_hi$Anno)) # Find relative to main fragment
logi_peak3_3 <- pac_filt_hi$Anno$Size <= 24 # Find relative to main fragment
logi_peak3a <- rowSums(cbind(logi_peak3_1, logi_peak3_2, logi_peak3_3)) == 3

pac_filt_hi$Anno$rRNA_peaks <- ifelse(logi_peak1, "peak1", 
                                   ifelse(logi_peak2, "peak2", 
                                           ifelse(logi_peak3a, "peak3a", 
                                                  ifelse(logi_peak4, "peak4", "other"))))

#--------
# Generate new PAC with only Peak fragments and plot errorbars:
pac_peaks <- PAC_filter(pac_filt_hi, anno_target=list("rRNA_peaks", c("peak1", "peak2", "peak3a", "peak4")))

dt_wide <- aggregate(pac_peaks$norm$cpm, list(pac_peaks$Anno$rRNA_peaks), sum)
nam_peaks <-  dt_wide[,1]
dt_wide <- as.data.frame(t(dt_wide[,-1]))
colnames(dt_wide) <- nam_peaks 
dt_wide$treatment <- pac_peaks$Pheno$treatment
dt_long<- reshape2::melt(dt_wide, id.vars="treatment")
targt <- c("DMSO_60", "BMH21_60", "BMH21_12")
dt_long$treatment <- factor(dt_long$treatment, levels=targt)

library(ggplot2)
ggplot(dt_long, aes(x=treatment, y=value, fill=treatment)) +
   							stat_summary(geom = "errorbar", fun.data = mean_se, position = "dodge") +
							stat_summary(geom = "bar", fun = mean, position = "dodge") +
							geom_jitter(col = "black", size=4, position=position_jitter(w=0.2, h=0.2)) +
                            facet_grid(. ~ variable, scales="free", space="free") +
							scale_fill_manual(values=c("#094A6B", "grey", "blue"))+
							scale_color_manual(values=c("black", "black", "black"))+
							theme_bw()+
							theme(legend.position="none", axis.line.x =element_blank())


dt_splt <-  split(dt_long, dt_long$variable)


#--------
# Non-parametric test

# First all peaks "DMSO_60" vs "BMH21_60" two sided
lapply(dt_splt, function(x){
  filt <-  x$treatment %in% c("BMH21_60", "DMSO_60")
  wilcox.test(x$value[filt]~x$treatment[filt])})  

# Then amplification (sided; less) from "BMH21_60" to "BMH21_12"
  peak1_dt <- dt_splt$peak1
  peak1_dt <- peak1_dt[peak1_dt$treatment %in% c( "BMH21_60", "BMH21_12"),]
  peak1_dt$treatment <- factor(peak1_dt$treatment, levels=c("BMH21_12","BMH21_60"))
  wilcox.test(peak1_dt$value~peak1_dt$treatment, alternative="less") 

# Then amplification (sided; less) from "BMH21_60" to "BMH21_12"
  peak4_dt <- dt_splt$peak4
  peak4_dt <- peak4_dt[peak4_dt$treatment %in% c( "BMH21_60", "BMH21_12"),]
  peak4_dt$treatment <- factor(peak4_dt$treatment, levels=c("BMH21_12","BMH21_60"))
  wilcox.test(peak4_dt$value~peak4_dt$treatment, alternative="greater")


  
#-------------------------------------------------------------------------------------
# Coverage plot of Peak 1 5' ETS rRF using Tong et al. 2020 dataset  (Fig 8K) 
#-------------------------------------------------------------------------------------

#--------
# Load previously saved filtered Tong et al. PAC object (section 2)
load("<your path to save folder/pac_filt_Tong2020.Rdata>")

#--------
# Remove corrupted cell lines and filter on pure human fragments (keep long fragments)  
pac_filt  <- PAC_filter(pac_filt, size = c(16,70), anno_target=list("pure", "pure_human"))
sampl <- as.character(unique(pac_filt$Pheno$cell_name))
sampl <- sampl[!sampl %in%  c("SCC-154", "SCC-4")]
pac_filt <- PAC_filter(pac_filt, pheno_target=list("cell_name", sampl))

#--------
# Re-normalize with pure human 
pac_filt <- pac_filt[!names(pac_filt)=="norm"]
pac_filt <- pac_filt[!names(pac_filt)=="summary"]
pac_filt <- PAC_norm(pac_filt, norm = "cpm")
pac_filt <- PAC_summary(pac_filt, norm="cpm", pheno_target=list("type"))

#--------
# Run PAC_mapper and PAC_covplot with same Supplementary_file_2.fa rRNA reference 
ref <- "<your path to Supplementary_file_2.fa>"
map_object <- PAC_mapper(pac_filt, ref=ref, N_up = "NNN", N_down = "NNN", mapper="reanno", mismatches=0, threads=8, report_string=FALSE)
out_cov <- PAC_covplot(pac_filt, map=map_object, summary_target=list("cpmMeans_type"), xseq = FALSE)
out_cov[2]


#-------------------------------------------------------------------------------------
# Clinical relevance for 5' ETS rRF in cervical cancer Xu et al dataset (Fig 8L) 
#-------------------------------------------------------------------------------------

# These scripts were used to generate the analysis reported in Fig 8L.
# We used a dataset originally published in:
#
#    Xu J, Zou J, Wu L, Lu W. Transcriptome analysis uncovers the diagnostic
#    value of miR-192-5p/HNF1A-AS1/VIL1 panel in cervical adenocarcinoma. Sci
#    Rep 2020 Oct 6;10(1):16584. PMID: 33024199
#
#    Xu J, Zhang Y, Huang Y, Dong X et al. circEYA1 Functions as a Sponge of
#    miR-582-3p to Suppress Cervical Adenocarcinoma Tumorigenesis via
#    Upregulating CXCL14. Mol Ther Nucleic Acids 2020 Dec 4;22:1176-1190. PMID:
#    33312754
#
# SRA project: SRP249688 
# ENA fastq download: https://www.ebi.ac.uk/ena/browser/view/PRJNA607023) 


# IMPORTANT, the Xu et al 2020 dataset is composed of both small RNA and long
# RNA fastq file. We only analyzed the sRNA (miRNA-seq) fastq files.

# IMPORTANT, this dataset originates from HiSeq X Ten sequencing using paired
# end 150+150 nt reads. Since the 2nd read is unnecessary for sRNA sequencing we
# did not include the paired 2nd read fastq files. Since 150 nt is well enough
# to detect the adaptor sequence, we used 'hard_rm' option in the 'adapt_3_set'
# input in the make_counts function.

#--------
# PAC generation
input <- "<your path to downloaded 1st read sRNA fastq files>"
pheno_path  = "<path to unpacked: Xu_pheno.csv>" # In Supplementary_file_S3.zip


parse <- list(polyG=c(type="hard_trim", min=20, mismatch=0.1),
                        adapt_3_set=c(type="hard_rm", min=10, mismatch=0.1),
                        adapt_3="AGATCGGAAGAGCACACGTCTGAACTCCA",
                        seq_range=c(min=14, max=70),
                        quality=c(threshold=20, percent=0.8),
                        indels=TRUE, concat=12, check_mem=TRUE)

count_list <- make_counts(input=input, type = "fastq", trimming="seqpac", 
                          parse=parse, threads=12, save_temp = TRUE)
anno <- make_anno(count_list, type = "counts", stat = TRUE)
pheno <- make_pheno(pheno_input=pheno_path,
                    counts=count_list$counts,
                    progress_report=count_list$progress_report)
pac  <- make_PAC(pheno_input=pheno, anno_input=anno, counts_input=count_list$counts)
table(pac$Pheno$tissue_type)
table(pac$Anno$Size)

#--------
# Filters and normalization 

pac_filt  <- PAC_filter(pac, size = c(16,70), threshold=3, coverage=50, pheno_target=list("tissue_type", c("cervical_tumor", "normal_cervical")), stat=TRUE)
pac_filt  <- PAC_norm(pac_filt, norm = "cpm")
pac_filt  <- PAC_norm(pac_filt, norm = "vst")
pac_filt  <- PAC_summary(pac_filt, norm = "cpm", pheno_target=list("tissue_type"))

nbias_out <- PAC_nbias(pac_filt, summary_target= list("cpmMeans_tissue_type", c("cervical_tumor", "normal_cervical")))
cowplot::plot_grid(plotlist=nbias_out$Histograms)
# Note, possible tRNA half peak in tumor samples. 


#--------
# Plot errorbars and non-parametric test (Fig 8L)  
ref <- "<your path to Supplementary_file_2.fa>"
map_object <- PAC_mapper(pac_filt, ref=ref, N_up = "NNN", N_down = "NNN", mapper="reanno", mismatches=0, threads=8, report_string=FALSE)

logi_peak1<- rownames(pac_filt$Anno) %in% rownames(map_object[[2]]$Alignments)
pac_filt$Anno$rRNA_peak_1 <- ifelse(logi_peak1, "peak1", "not_peak")
pac_peaks <- PAC_filter(pac_filt, anno_target=list("rRNA_peak_1", c("peak1")))

dt_wide <- aggregate(pac_peaks$norm$cpm, list(pac_peaks$Anno$rRNA_peak_1), sum)
nam_peaks <-  dt_wide[,1]
dt_wide <- as.data.frame(t(dt_wide[,-1]))
colnames(dt_wide) <- nam_peaks 
dt_wide <- cbind(dt_wide, pac_peaks$Pheno$tissue_type)
colnames(dt_wide)[2] <- "type"
library(ggplot2)
ggplot(dt_wide, aes(x=type, y=peak1, fill=type)) +
   							stat_summary(geom = "errorbar", fun.data = mean_se, position = "dodge") +
							stat_summary(geom = "bar", fun = mean, position = "dodge") +
							geom_jitter(col = "black", size=4, position=position_jitter(w=0.2, h=0.2)) +
							scale_fill_manual(values=c("#094A6B", "grey", "blue"))+
							scale_color_manual(values=c("black", "black", "black"))+
							theme_bw()+
							theme(legend.position="none", axis.line.x =element_blank())

wilcox.test(dt_wide$peak1~dt_wide$type) 


#######################################################################################
#####  4. 5' ETS rRF is missing in Snoek et al 2018 dataset (unpublished) 
####################################################################################### 

# These scripts were not represented by any figures in the main text. We
# provided it for future explorations of the 5' ETS rRF, as we were unable to
# confirm this rRF in this dataset, despite finding it in 3 other unrelated
# datasets. We speculate that this is due to library preparation kit used, since
# the 3 datasets confirming the 5' ETS rRF were prepared using the NEBNExt Small
# RNA library prep kit, while Snoek et al 2018 used the Illumina TrueSeq Small
# RNA Library Preparation kit.
#
# The dataset was originally published in:
#
#   Snoek BC, Verlaat W, Babion I, Novianti PW et al. Genome-wide microRNA analysis
#   of HPV-positive self-samples yields novel triage markers for early detection of
#   cervical cancer. Int J Cancer 2019 Jan 15;144(2):372-379. PMID: 30192375
#
# SRA project: PRJNA413777
# ENA fastq download: https://www.ebi.ac.uk/ena/browser/view/PRJNA413777) 


#--------
# PAC generation
input <- "<your path to downloaded Snoek et al fastq files>"
pheno_path  = "<path to unpacked: Snoek_pheno.csv>" # In Supplementary_file_S3.zip

# They used Illumina TrueSeq; Average size 50 nt (use hard_trim)
parse <- list(polyG=c(type="hard_trim", min=20, mismatch=0.1),
                        adapt_3_set=c(type="hard_trim", min=10, mismatch=0.1),
                        adapt_3="TGGAATTCTCGGGTGCCAAGGAACTCCAGTCAC",
                        seq_range=c(min=14, max=70),
                        quality=c(threshold=20, percent=0.8),
                        indels=TRUE, concat=12, check_mem=TRUE)

count_list <- make_counts(input=input, type = "fastq", trimming="seqpac", 
                          parse=parse, threads=12, save_temp = TRUE)
anno <- make_anno(count_list, type = "counts", stat = TRUE)
pheno <- make_pheno(pheno_input=pheno_path,
                    counts=count_list$counts,
                    progress_report=count_list$progress_report)
pac  <- make_PAC(pheno_input=pheno, anno_input=anno, counts_input=count_list$counts)

#--------
# Filters and normalization: 
pac_filt  <- PAC_filter(pac, size = c(14,70), threshold=3, coverage=50, pheno_target=list("histology", c("CIN3", "normal")), stat=TRUE)
pac_filt  <- PAC_norm(pac_filt, norm = "cpm")
pac_filt  <- PAC_norm(pac_filt, norm = "vst")
pac_filt  <- PAC_summary(pac_filt, norm = "cpm", pheno_target=list("histology"))


#--------
# Map and generate coverage plots same reference as the other datasets: 
ref <- "<your path to Supplementary_file_2.fa>"
map_object <- PAC_mapper(pac_filt, ref=ref, N_up = "NNN", N_down = "NNN", mapper="reanno", mismatches=0, threads=8, report_string=FALSE)

out_covp <- PAC_covplot(pac_filt, map=map_object, summary_target= list("cpmMeans_histology", c("CIN3", "normal")), xseq = FALSE)
out_covp[[1]] # Much variability, and different peaks than NEBNExt kit
out_covp[[2]] # Very few fragments in Peak 1.
```
